# The acute phase response in bats (*Carollia perspicillata*) varied with the time and dose of the immune-challenge

**DOI:** 10.1101/2022.05.18.492341

**Authors:** Matheus F. Viola, L. Gerardo Herrera M., Ariovaldo P. da Cruz-Neto

## Abstract

The acute phase response (APR) is a core component of the innate immune response and represents the first line of immune defense used in response to infections. Although several studies with vertebrates reported fever, decrease in food intake and body mass, as well as increase in neutrophil/lymphocyte ratio and total white blood cell count after lipopolysaccharide (LPS) inoculation there was great variability in the magnitude of these responses. Some of these differences might reflect, to some extent, differences in time of endotoxin inoculation (during activity or resting periods) and dose; however, no study in the literature has evaluated the synergistic effect of these two factors in vertebrates. Therefore, our study tested the interplay between LPS dose and time of injection on selected physiological (fever and increase in total white blood cell count and neutrophil/lymphocyte ratio), and behavioral (food intake) components of APR using a Neotropical fruit-eating bat (*Carollia perspicillata*) as a model organism. We predicted that LPS would trigger a dose- and time-dependent response on APR components. APR components were assessed in resting and activity periods after injection of three doses of LPS (5, 10 and 15 mg/kg LPS). The results showed that LPS-evoked changes in skin temperature, food intake, neutrophil lymphocyte ratio depend markedly on the LPS dose and/or time that LPS is administered.

## Introduction

Animals are constantly exposed to a myriad of pathogens, and their immune system plays a pivotal role in mitigating the potential deleterious effects of such exposure (Acevedo-Whitehouse and Duffus 2009). From a methodological perspective, ecoimmunological studies ultimately have attempted to use some metrics that portrait immunocompetence, that is, the ability of the immune system to fight pathogens (Demas et al. 2011; Schoenle et al. 2018). The acute phase response (APR) is a core component of the innate immune response, and its quantification is widely used to assess the immunocompetence of different species of animals (Merlo et al. 2016; Roy et al. 2016; Ramirez-Otarola et al. 2018). APR is the first line of induced defense used by all animals in response to infections, and the most common technique employed to activate it in experimental studies is the central or peripherical administration of lipopolysaccharide (LPS - Rudaya et al. 2005; Cray et al. 2009; Sampath 2018).

LPS is an immunogenic component of the outer membrane of gram-negative bacteria that causes the release of proinflammatory cytokines, which in turn trigger a suite of behavioral (decrease activity, anorexia, adipsia), and physiological responses (body temperature changes, increase in total white blood cell count and activation of the hypothalamic–pituitary–adrenal axis - Lochmiller and Deerenberg 2000; Sampath 2018). APR accelerates pathogen elimination and enhances the activation of the adaptive immune system, thus conferring an immediate benefit in controlling infection (Cray et al. 2009). APR, however, is thought to have a high cost/benefit ratio because its protective value is lower when compared to adaptive constitutive responses, such as those mediated by leukocytes, and the energetic cost of its activation is the highest of all immune system defenses (Martin et al. 2007; Hasselquist and Nilsson 2012). The overall energetic cost of LPS-induced APR and/or the magnitude of its behavioral and physiological components, have been measured in several taxons. Although several studies report an increase in metabolic rate, and a decrease in food intake and body mass after LPS inoculation, there was a great variability in the magnitude of these responses among species (Volkoff and Peter 2004; Burness et al. 2010; Llewellyn et al. 2011; Marais et al. 2011; MacDonald et al. 2012; King and Swanson 2013). On the other hand, not all studies have reported that LPS triggers fever, increases metabolic rate, activates the HPA axis (hypothalamic–pituitary–adrenal), or increases the total white blood cell count (Weber et al. 2005; Copeland et al. 2005; Krams et al. 2012; Stockmaier et al. 2015; Rakus et al. 2017; Lind et al. 2020; Amaral-Silva et al. 2021). Some of these differences certainly reflect species-specific differences and they are contingent on the specific context of the study. However, some of these variations might also reflect differences in time and dose of LPS injections.

For example, LPS triggers a decrease in food intake when injected during the activity period of *Mus musculus* but not when injected during the resting period (Mathias et al. 2000). This rodent exhibits a daily rhythm of leukocytes, peaking in the circulation during the resting period, whereas recruitment of leukocytes to tissues is higher during the active period after LPS injection (Scheiermann et al. 2012, 2013). Activation of HPA axis in toads followed LPS injection is more pronounced during activity period compared with resting period (Titon et al. 2022). In birds, LPS injection triggers fever response during the resting period (*Melospiza melodia*; Adelman et al. 2010; *Passer domesticus*; Coon et al. 2011; *Taeniopygia guttata*; Skold-Chiriac et al. 2015), with a brief hypothermic response during the activity period (*Taeniopygia guttata*; Owen-Ashley et al. 2006; *Zonotrichia leucophrys gambelii*; Burness et al. 2010). LPS injection set off a long-lasting fever response in *Columba livia* during the activity period (Nomoto 1996), while in *Mus musculus* it set off a fever response during the resting period with a brief hypothermic response followed by fever during the activity period (Morrow and Opp 2005).

Studies carried out in birds and small rodents show that a decrease of food intake and body mass is dose-dependent, with highest doses resulting in a higher decrease of food intake and body mass (0.05 to 3 mg/kg LPS; Kozak et al. 1994; 0.3 to 5 µg/kg LPS; Haba et al. 2012; 0.001 to 1 mg/kg LPS; Skold-Chiriac et al. 2015). In goldfish (Carassius auratus) there is a dose dependent effect of LPS on food intake, with higher doses leading to greater decrease of food intake (10 to 250 ng/g LPS; Volkoff and Peter 2004). Rabbits challenged with higher LPS doses show a higher increase in total white blood cell count and increased neutrophilia and lymphopenia (0.01 to 1 µg/kg LPS; Kimura et al. 1994). Toads injected with higher LPS dose show a longer-lasting behavior fever response (0.2 to 2 mg/kg LPS; Bicego et al. 2002). Turtles injected with lower LPS dose (0.0025 mg/kg LPS) show hypothermic behavior response, whereas when injected with a higher dose show behavior fever response (0.025 mg/kg LPS; Do Amaral et al. 2002). Higher LPS doses triggers a longer lasting fever response and higher peak of fever in pekin ducks (*Anas platyrhynchos domesticus*; 0.001 - 1 mg/kg LPS; Maloney and Gray 1998), while in *Galus galus* lower LPS doses (0.002 to 0.01mg/kg LPS) results in fever response, and a higher LPS dose (0.1 mg/kg LPS) results in hypothermic response followed by fever (Dantonio et al. 2016; Amaral-Silva et al. 2020). A higher LPS dose results in a longer lasting hypothermic response in Zebra finch (*Taeniopygia guttata*; 0.001 – 1 mg/kg LPS; Skold-Chiriac et al. 2015). Injections of higher LPS doses set off a long last fever response in small rodents, and in some case, this response is preceded for a brief hypothermic response after higher dose (0.003 to 1 mg/kg LPS; Romanovsky et al. 1996a, 1996b; Steiner et al. 2009; 5 to 18 mg/kg LPS; Liu et al. 2012). Analyzes of bats’ immune response repertoire are fundamental to understand why this order hosts a wide variety of pathogens without showing clear signs of developing disease in most cases (O’Shea et al. 2014; Moratelli and Calisher 2015; Kacprzyk et al. 2017). During their active period, bats increase their exposure to pathogens while foraging (Wong et al. 2007; Patz et al. 2008; Kuzmin et al. 2011), whereas during resting, aggregation behavior within their shelters and grooming increases the potential for transmission of infectious agents (Mühldorfer 2013; White and Razgour 2020). In this sense, the activation of APR in bats might occur during the activity period or during the resting period, and the magnitude of this response should vary with pathogenic load. Studies that have analyzed APR in bats examined only certain components of the APR during specific periods and using a single LPS dose. For example, LPS injections triggered fever in *C. perspicillata* (2.8 mg/kg LPS; Cabrera-Martinez et al. 2019), *Myotis vivesi* (1.75 mg/kg LPS; Otálora-Ardila et al. 2016) and *Artibeus lituratus* (2 mg/kg LPS; Guerrero-Chacón et al. 2018) when administered during the resting period, but no change on body temperature was observed when administered during the activity period in *C. perspicillata* (3 mg/kg LPS; Melhado et al. 2020) and *Molossus molossus* (4.5 mg/kg LPS; Stockmaier et al. 2015). Some studies report leukocytosis after LPS injections during the resting (2 mg/kg LPS; Schneeberger et al. 2013; Weise et al. 2017) or activity periods (5 mg/kg LPS; Stockmaier et al. 2018), while others show no evidence of leukocytosis during the activity (Stockmaier et al. 2015; Melhado et al. 2020) or resting periods (Guerrero-Chacón et al. 2018; Cabrera-Martinez et al. 2019). Leukocytosis in bats is associated with increased circulating neutrophils (Weise et al. 2017), whereas the N/L ratio increased after LPS injection with similar doses of LPS during the active or resting phases (Stockmaier et al. 2018; Cabrera-Martinez et al. 2019; Melhado et al. 2020). Although most of these studies reported body mass loss 24 hours after LPS injections and assumed that it is partly due to anorexia, not all examined the effect of LPS on food intake rate (Schneeberger et al. 2013; Stockmaier et al. 2015, 2018; Guerrero-Chacón et al. 2018).

In this study, we tested the interplay between LPS dose and time of injection (during activity and resting periods) on selected physiological (body temperature change, body mass change, total white blood cell count, and neutrophil/lymphocyte ratio) and behavioral (food intake) components of the APR using a Neotropical fruit-eating bat (*Carollia perspicillata*) as a model organism. We predicted that LPS would trigger a dose-dependent and/or period-dependent response on bat’s APR components. We expected that the highest LPS doses would result in a higher decrease of food intake and body mass loss, as well as a higher increase of the neutrophil/lymphocyte ratio and the leukocyte count, and that these responses would be more pronounced when LPS was injected at the beginning of the activity period. We further expected that LPS would trigger a dose-dependent change on body temperature during the resting and activity periods, with highest LPS doses resulting in an increase on magnitude and/or duration of fever response during the resting period. However, given the absence of thermoregulatory response observed in bats during the activity period (Stockmaier et al. 2015; Melhado et al. 2020), and the variety of thermoregulatory responses observed in small rodents and birds during this period (Morrow and Opp 2005; Owen-Ashley et al. 2006; Amaral-Silva et al. 2020) we made no predictions about the direction of body temperature change (e.g., fever or hypothermia).

## Material and methods

### Capture and maintenance

Non-reproductive adults of *C. perspicillata* were captured between June 2018 and July 2019 in 3 municipalities at São Paulo State, Southeastern Brazil: Edmundo Navarro de Andrade State Forest, located in Rio Claro (22°25’54,2’’S 47°32’11,1’’W), forest remnants of the Federal University of São Carlos - UFSCAR, located in São Carlos (23°21’32,5’’S 46°15’15,1’’W), and RPPN Rio dos Pilões located in Santa Isabel (21°59’6,2’’S 47°52’55,3’’W). Bats were captured with mist nets and transported to an outdoor cage (3 × 3 × 3 m), exposed to natural conditions of photoperiod and temperature (22 ± 1°C SD; CEAPLA/IGCE/UNESP) at Universidade Estadual Paulista at Rio Claro. The bats were fed papaya and bananas for five days before the immune challenge. Permits to capture and housing bats were issued by the Instituto Chico Mendes de Conservação da Biodiversidade (ICMBio, process number 66452-1). Ethical permits for this study were issued by the Animal ethics committee of the Universidade Estadual Paulista at Rio Claro (Authorization: n° 3381).

### Experimental conditions and immune challenge

The immune challenges were conducted in a climatic chamber (3 × 2 × 3 m), with controlled room temperature (27°C – thermoneutral zone of *C. perspicillata* - Cruz-Neto and Jones, 2006) and photoperiod (12/12 hours). Lights were switched on at 06:00 am and switched off at 06:00 pm. Bats were transferred to the climatic chamber one day before immune challenge for acclimatization, and they were kept in individual cages (1 × 1 × 0.5 m). After acclimatization, 24 bats were assigned to the immune challenge at 06:00 pm, and 24 bats were assigned to the immune challenge at 08:00 am using three LPS doses and one control group. Six bats of each injection period group (night- and day-time injections) were randomly assigned to either the LPS groups or the control group. Bats were injected subcutaneously into the scapula with 5 mg/kg, 10 mg/kg or 15 mg/kg of LPS (LPS L2630; Sigma-Aldrich) diluted in 50 ul of phosphate-buffered saline (PBS; P4417, Sigma-Aldrich, USA), while control groups were injected only with 50 ul of PBS. In this way, we conducted 4 experimental trails for day- and nighttime injections, respectively. The experimental trials were conducted on the climatic chamber with 6 individuals at a time, and the bats were assigned to only one period and dose treatments. The range of LPS doses represent the maximum dose used in previous studies with bats (5 mg/kg LPS - Stockmaier et al. 2015; 2018) and doses at least two to three times that value.

### Food intake and body mass variation

Before and after injections, bats were fed ad libitum with mango nectar (> 95% of the composition in the form of simple sugars - Serigy®) supplemented with 4 mg/l of hydrolyzed casein (Sigma Aldrich®). A known amount of nectar was placed in a feeding device at 06:00 pm and was weighed at 08:00 am to measure food intake.

The amount of food given exceeded daily consumption based on preliminary observations. We also placed a feeding device in the climactic chamber with a known amount of nectar to measure evaporative loss. We assessed food intake changes in relative terms as: food intake change (ΔFI) = (food intake after injections – food intake before injections) / (food intake before injections). Bats were weighed to the nearest 0.1 g (Ohaus Precision Balance, USA) before and after injections. Body mass of bats injected during the activity period was measured at 06:00 pm and 08:00 am before injection, right before the injection at 06:00 pm, and after the injection at 08:00 am and 06:00 pm. Body mass of bats injected during the resting period was measured around 08:00 am and 06:00 pm before injection, right before the injection at 08:00 am, and after the injection at 06:00 pm and 08:00 am. We assessed the body mass changes in relative terms as: body mass change (ΔMb) = (mean body mass after injections mean body mass before injections) / (mean body mass before injections), (see Supplementary Material 1; Fig. 1 and 2).

**Fig. 1.**
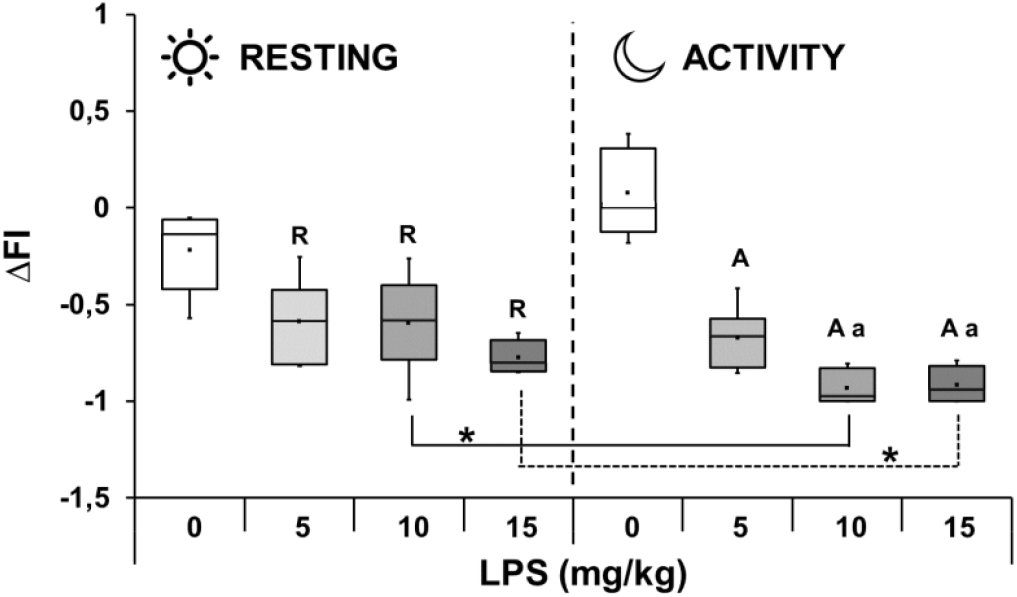
Food intake change (ΔFI) after PBS and LPS injections (5, 10 and 15 mg/kg LPS) during resting and activity periods in *Carollia perspicillata*. (**R**) represents significant differences (P ≤ 0.05) between PBS and LPS injected groups during the resting period. N = 5 for PBS injected groups. N = 6 for all other groups. (**A**) represents significant differences (P ≤ 0.05) between PBS and LPS injected groups during the activity period. N = 6 for all groups. (**a**) represents significant differences (P ≤ 0.05) between 5 and 10 mg/kg LPS, and between 5 and 15mg/kg LPS injected groups during the activity period. (**\***) represents significant differences (P ≤ 0.05) between LPS injections groups during the resting and activity periods

**Fig. 2.**
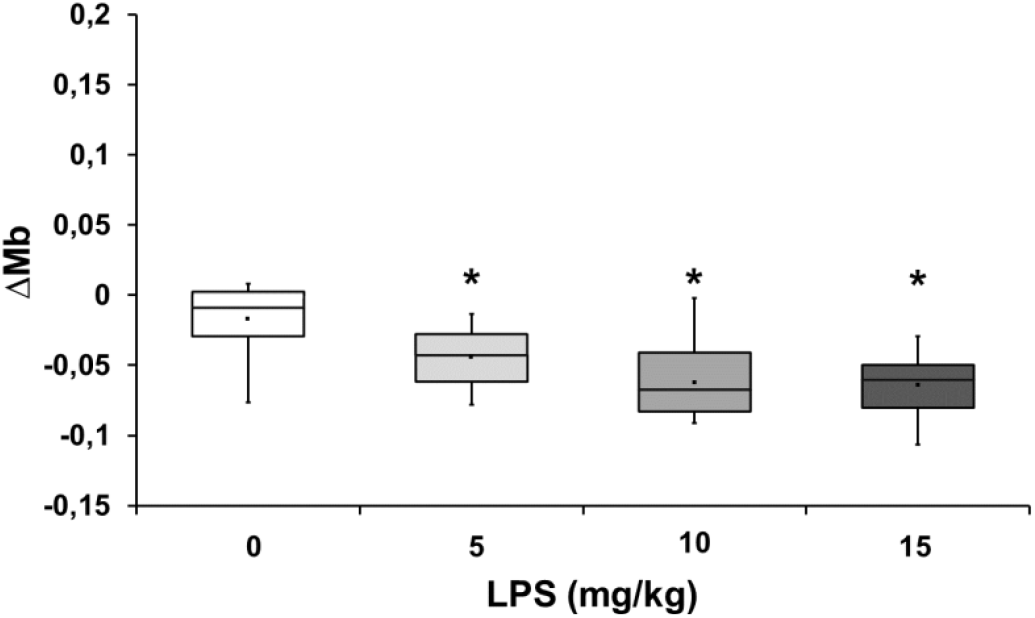
Mean body mass change (ΔMb) after PBS and LPS injections (5, 10 and 15 mg/kg LPS dose) regardless of injection period in *Carollia perspicillata*. (**\***) represents significant differences (P ≤ 0.05) between the PBS and LPS injected groups. N = 6 for all groups

### Total white blood cells count (WBCs) and Neutrophil / Lymphocyte Ratio (N/L)

We collected ∼10–15 μl of blood from the propatagial vein 24 hours before and after injections at around 06:00 pm (groups injected during the activity period) or 08:00 am (groups injected during the resting period) and prepared two blood smears to examine WBCs. We counted WBCs in 20 field views of each blood smear at ×400 magnification in a microscope (Melhado et al. 2020; Moreno et al. 2021). In addition, we counted neutrophil and lymphocyte at ×1000 magnification in two replicates of sets of 100 WBCs on each blood smear to calculate the N/L ratio (Paksuz et al. 2009; Melhado et al. 2020). We assessed WBCs and N/L changes in relative terms as: WBCs changes (ΔWBC) = (WBCs after injections – WBCs before injections)/ (WBCs before injections); and N/L changes (ΔN/L) = (N/L after injections – N/L before injections)/ (N/L before injections).

### Body temperature variation

We measured skin temperature as an approximation of body temperature (Williams et al. 2009) using Sub-Cue Temperature Transmitters (2.71 ± 0.05 g; Canadian Analytical Technologies, Calgary, Canada) attached in the skin of bats on the scapular region 24 hours before injections. Measurement of skin temperature is considered a good estimator of body temperature in bats (Otálora-Ardila et al. 2016; Melhado et al. 2020). Body temperature was recorded every hour from 24 hours before to 24 hours after injections, starting at 06:00 pm in groups injected during the activity period, and at 06:00 am in groups injected during the resting period. We assessed skin temperature change 12 hours after injections in absolute terms (ΔTb) by subtracting hourly skin temperature after injections (STAhourly) from the respective hourly skin temperature before injections (STBhourly; see Supplementary Material 1; Fig. 3 and 4). We used the values of temperature from the calibration curves provided by the manufacturer before checking their readings against values obtained simultaneously with a thermometer and with eight the transmitters placed in the climate chamber at different temperatures. Values obtained with the calibration curve and direct readings differed by 0.22°C ± 0.11 (mean ± SD).

**Fig. 3.**
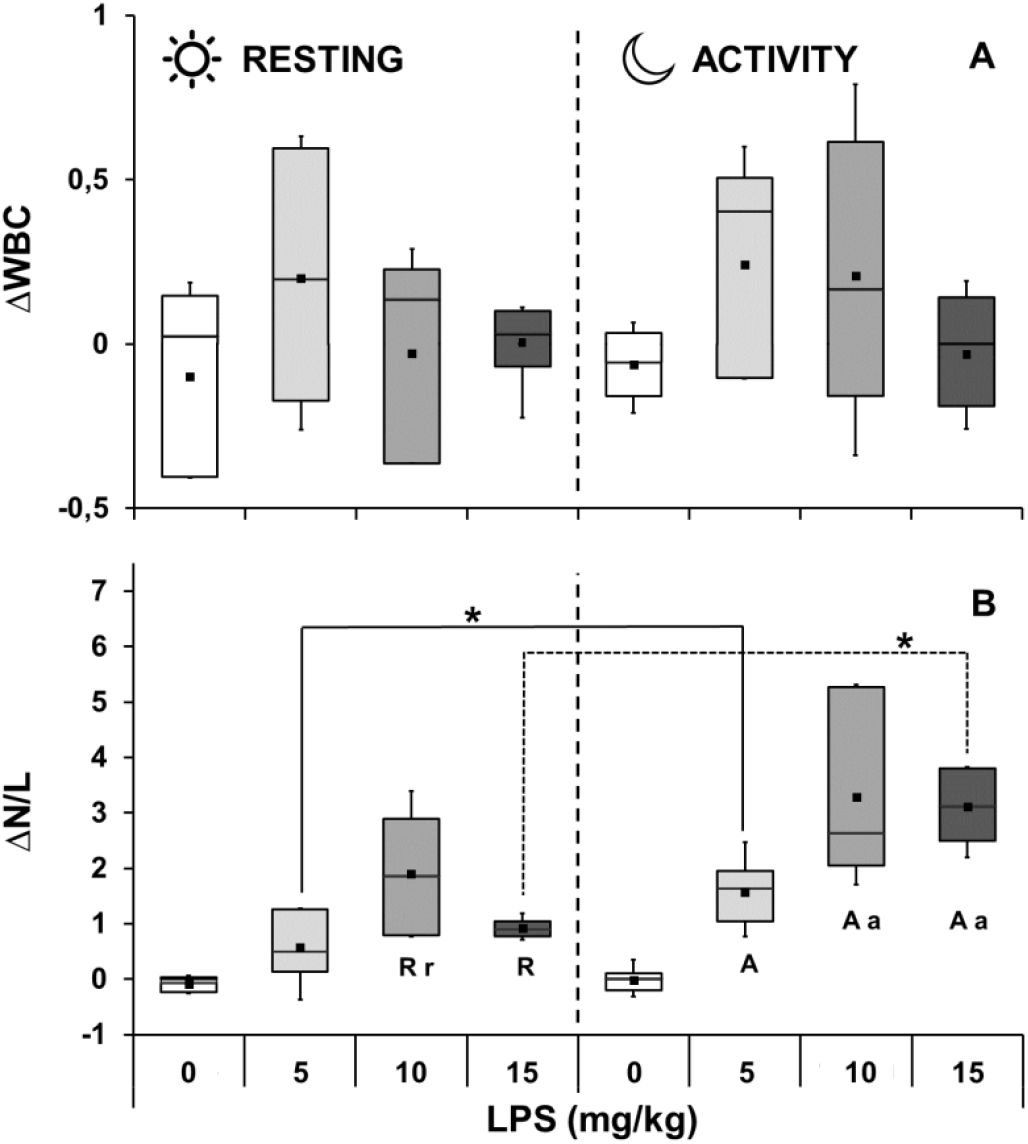
Total leucocyte count change (ΔWBC) and Neutrophil/Lymphocyte ratio change (ΔN/L) after PBS and LPS injections (5, 10 and 15 mg/kg LPS) during resting and activity periods in *Carollia perspicillata*. **A:** ΔWBC after injections during the resting and activity periods. N = 6 for all groups. **B:** ΔN/L after injections during the resting and activity periods. N = 6 for all groups. (**R**) represents significant differences (P ≤ 0.05) between PBS and LPS injected groups during the resting period. (**r**) represents significant differences (P ≤ 0.05) between 5 and 10mg/kg LPS injected groups during the resting period. (**A**) represents significant differences (P ≤ 0.05) between PBS and LPS injected groups during the activity period. (**a**) represents significant differences (P ≤ 0.05) between 5 and 10mg/kg LPS, and between 5 and 15 mg/kg LPS injected groups during the activity period. (*) represents significant differences (P ≤ 0.05) between LPS injected groups during the resting and activity periods

**Fig. 4.**
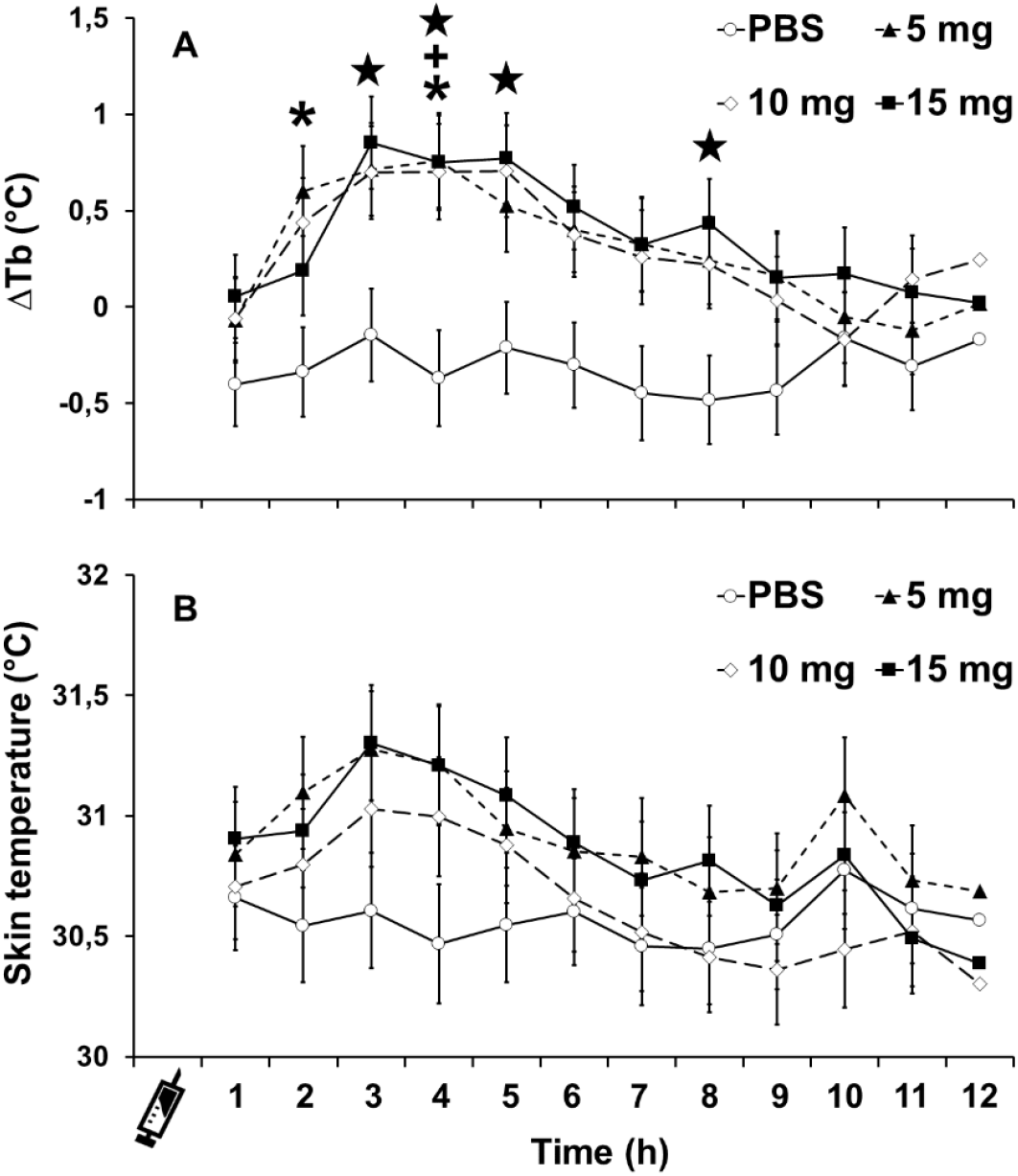
Hourly skin temperature changes (ΔTb) and hourly skin temperature (°C) after PBS and LPS injections (5, 10 and 15 mg/kg LPS) in *Carollia perspicillata*. **A:** Hourly skin temperature changes (ΔTb) between and within PBS and LPS injected groups. (*) represents significant difference (P ≤ 0.05) between 5mg/kg LPS and PBS injected groups. (**+**) represents significant differences (P ≤ 0.05) between 10mg/kg LPS and PBS injected groups. (★) represents significant difference (P ≤ 0.05) between 15mg/kg LPS and PBS injected groups. **B:** Hourly skin temperature (°C) after PBS and LPS injected groups (5, 10 and 15 mg/kg LPS). N = 6 for all groups. All figures show mean ± standard error

### Data analysis

A two-way ANOVA was used to test for the interaction effects of dose (PBS, 5 mg/kg LPS, 10 mg/kg LPS, and 15 mg/kg LPS) and time of injection (resting and activity periods) on food intake, body mass change, WBC and N/L ratio. If the two-way interaction term of these analysis were significant (dose x time of injection), the simple main effects of dose were analyzed with a one-way ANOVA for each time of injection separately, followed, when applicable, by a Tukey HSD post-hoc test for pairwise means comparisons. If the effects of dose were significant in both time of injection, we tested for dose-effect differences between time of injection (simple effect of time of injections) with a set of independent t-test. Main effects of two-way ANOVA were interpreted in the absence of statistically significant interactions (Maxwell and Delaney 2004; Kiernan 2014) by a Tukey HSD post-hoc test for pairwise comparisons when applicable (Christopoulos et al. 2019; Williams et al. 2019; see Supplementary Material 1; Fig. 5). Three-way mixed ANOVA (2 Between subject factors / 1Within subject factor) were used to test for the interaction effects of dose (PBS, 5 mg/kg LPS, 10 mg/kg LPS, and 15 mg/kg LPS) and time of injection (resting or activity) as between subject factors, and time (ΔTb for 12 hours) as within subject factors on skin temperature change (ΔTb). When three-way interaction or two-way interaction were significant, simple main effect of factors (dose, time of injections and time) were tested by a Bonferroni post-hoc test for pairwise means comparisons (Rygula et al. 2006; Csabafi et al. 2018; Cohen et al. 2020; Guan et al. 2020; Sun et al. 2021). Main effect Three-way mixed ANOVA were interpreted in the absence of statistically significant interactions (Maxwell and Delaney 2004; Kiernan 2014) by Tukey HSD or Bonferroni post-hoc test for pairwise comparisons when applicable. All variables were checked for normality (Shapiro-Wilk test) and homogeneity (Levene’s test), which validated the use of parametric tests. In cases where one of the assumptions were violated, data were transformed to meet the assumptions of normality and homogeneity. Accordingly, ΔWBC was transformed to cubic root, and log transformation was applied on ΔN/L. Studentized residuals test were used to detect significant outliers (± 3 standard deviation), which were then removed from the analysis. Sphericity was assessed with Mauchly’s test, and the Greenhouse-Geisser correction was applied when necessary. All statistical analyses were performed in SPSS 26 for Windows (IBM Corp., Armonk, NY), and a fiducial level of 0.05 was adopted to determine the significance of all comparations. Before injections, there was no difference in food intake, mean body mass, total white blood cell count, neutrophil/lymphocyte or mean skin body temperature among localities where bats were collected (see Supplementary Material 1; Table 1).

## Results

### Food intake (ΔFI) and Body mass (ΔMb)

There were significant interaction effects of dose and time of injection on food intake change (F_3,38_ = 6.10, p = 0.002, partial η^2^ = 0.325). Food intake decreased when bats were injected during the day (F_3.19_ = 7.3, p = 0.002, partial η^2^ = 0.535; Fig. 1). In this case, all injected groups showed a mean decrease of food intake, but the magnitude of decrease was higher in the groups injected with 5, 10 and 15 mg/kg LPS doses compared to PBS injected ones (59, 60 and 77% respectively; p < 0.050; Fig. 1). No significant difference was found in mean decrease food intake among bats injected different LPS doses (p > 0.050). For bats injected during activity, injections also affected food intake (F_3.19_ = 53.8, p < 0.001, partial η^2^ = 0.89; Fig. 1). In this case, decreased mean food intake was higher for bats injected with 5, 10 and 15 mg/kg LPS doses when compared to PBS-injected bats (p < 0.050), and the magnitude of this decrease was higher for bats injected with of 10 or 15 mg/kg LPS doses than when injected with 5 mg/kg LPS dose (67, 93 and 92% respectively; p < 0.050; Fig. 1). However, no significant difference was observed in decrease mean food intake between bats injected with 10 or 15mg/kg LPS doses (p > 0.050). The changes in food intake were not significantly different between PBS-injected bats during the resting and activity periods (t_8_ = -2.12; p = 0.066), whereas the magnitude of decrease in mean food intake was more pronounced for bats injected with 10 and 15 mg/kg LPS doses during the activity period compared to the respective groups injected during the resting period (t_10_ = 3.12; p = 0.011; t_10_ = 2.72; p = 0.022 respectively; Fig. 1).

There was only a significant main effect of dose on mean body mass change (F_3.40_ = 11.28, p < 0.001, partial η^2^ = 0.458; Fig. 2). All injected groups showed a mean decrease of mean body mass, but the magnitude of the decrease was higher in bats injected with 5, 10 and 15 mg/kg LPS doses (4.4, 6.3 and 6.4% respectively; p < 0.050; Fig. 2) compared to those injected PBS (2%). No significant difference was found in mean body mass change among bats injected with LPS doses (p > 0.050).

### White blood cell (ΔWBC) and Neutrophil/Lymphocyte ratio(ΔN/L)

No significant dose-time of injection interaction (F_3.40_ = 0.48, p = 0.701, partial η^2^ = 0.034), nor for the main effects of dose (F_3.40_ = 0.880, p = 0.460, partial η^2^ = 0.062) and time of injection (F_1.40_ = 0.07, p = 0.799, partial η^2^ = 0.002) were observed for WBC change (Fig. 3A). On the other hand, there was a significant interaction effect of dose and time of injection on N/L change (F_3,40_ = 3.11, p = 0.037, partial η^2^ = 0.189). N/L change varied among doses when bats were injected during the resting (F_3,20_ = 13.46, p < 0.001, partial η^2^ = 0.669): mean N/L ratio increased at a higher rate for bats injected with 10 and 15 mg/kg LPS when compared to PBS-injected bats (189 and 91% respectively; p < 0.050), and the magnitude of this increase was higher for bats injected with of 10 mg/kg LPS than with 5 mg/kg LPS (p < 0.050; Fig. 3B). No significant difference was observed in N/L change between bats injected with 5, 10 or 15 mg/kg LPS doses (p > 0.050). For bats injected during the activity period, injection dose also affected mean N/L change (F_3,20_ = 42.40, p < 0.001, partial η^2^ = 0.864). In this case, mean N/L change was higher for bats injected with 5, 10 and 15 mg/kg LPS when compared to PBS-injected bats (157, 328 and 311% respectively; p < 0.050), and the magnitude of this increase was higher for bats injected with of 10 and 15 mg/kg LPS than with 5 mg/kg LPS (p < 0.050; Figure 3B). However, no significant difference was observed in N/L change between bats injected with 10 or 15 mg/kg LPS doses (p > 0.050). The mean N/L ratio changes were not significantly different between PBS-injected bats during the resting and activity periods (t _10_ = -0.59; p = 0.569). The magnitude of the increase in N/L ratio was more pronounced for bats injected with 5 and 15 mg/kg LPS doses during the activity period than those injected the same dose during the resting period (t_10_ = -2.54; p = 0.029; t_10_ = -9.94; p < 0.001 respectively; Fig. 3B). Bats injected with 10 mg/kg LPS during the activity period showed a higher increase of N/L ratio compared with those injected the same dose during the resting period, but this increase was not significant (t_10_ = -1.86; p = 0.093).

### Skin temperature (ΔTb)

There was a significant main effect of time (F_4.26_,_170.69_ = 1.26, P < 0.001, partial η^2^ = 0.864) and dose (F_3.40_ =3.62, P = 0.021, partial η^2^ = 0.213) on hourly skin temperature change, and the dose-time interaction (F_12.80,170.69_ = 1.81, P = 0.045, partial η^2^ = 0.120), indicating that that this change overtime depends on the dose, but no on the period of injection. Hourly skin temperature of bats injected with PBS did not change with time after injection (p > 0.404; Fig. 4A). Two hours after injections there was a significant increase in skin body temperature of 5 mg/kg LPS injected bats compared to the first hour (p < 0.001; Fig. 4A), whereas 10 and 15 mg/kg LPS injected bats increased their skin temperature three hours after injection compared to the first hour (p < 0.001; Fig. 4A). Skin temperature change of bats injected with PBS, 5, 10 and 15 mg/kg LPS was not significantly different 1 hour after injections (p > 0.993). Two hours after injections, only bats injected with 5 mg/kg LPS showed a significant increase in skin temperature compared with those injected PBS (0.6 °C; p < 0.021; Fig. 4A). Three hours after injections only bats injected with 15 mg/kg LPS dose showed a significative increase in skin temperature compared with bats injected PBS (0.9°C; p < 0.021; Fig. 4A). Four hours after injections, bats injected with 5, 10 and 15 mg/kg LPS dose showed a significative increase in skin temperature compared with PBS-injected bats (0.8, 0.7 and 0.8°C respectively; p < 0.050; Fig. 4A). Five hours after injections only bats injected with 15 mg/kg LPS showed a significant increase in skin temperature compared with PBS-injected bats (0.8°C; p < 0.050; Fig. 4A). No difference was found on skin temperature changes among bats that received LPS doses at any time point (p > 0.05). After the maximum skin temperature response, bats of all LPS injected groups steadily decrease their body temperature, converging to the body temperatures of PBS group (p > 0.050; Fig. 4A). Eight hours after injections bats injected with 15 mg/kg LPS showed a new significative increase in skin temperature compared with PBS (0.5°C; p < 0.05; Figure 4A), but this increase was not significantly different than the increase observed between 3 and 5 hours after injection (p < 0.05; Fig. 4A).

## Discussion

To our knowledge this is the first study investigating the interplay between dose and time of injection on components of APR in vertebrates. We expected that LPS would trigger a dose-dependent and/or time-dependent response on bat’s APR components. Our results indicate that some components of APR evaluated in this study were differently affected by dose (body mass variation and skin temperature) or by the interaction between dose- and time of injection (food intake and N/L ratio). Interestingly, even using LPS doses at least 3 times higher than those used in previous studies with birds and small rodents, total white blood cell count was not significantly affected regardless of the dose or period of injections.

Food intake decreased in a dose-dependent manner after injections during the activity period, but no after injections during the resting period. The acute phase response occurs immediately after pathogen infection, lasting approximately 12 to 24 hours (Demas and Nelson 2012). When LPS is administered during the day (resting period), bats experience the first hours of induced disease behavior (anorexia and reduced activity) at a time when there is no food intake and activity is reduced, but when LPS is administered at night bats experience the first hours of induced disease behavior at the time of feeding, when they are naturally more active (Weise et al. 2017). Food intake decreased from the first to the second night of the experiment regardless of injection treatment during the resting period, but this decrease was at least more than double for bats injected with LPS (59 - 77% decrease) than for bats injected with PBS (22% decrease). Despite the decrease in food intake observed in the PBS-injected group during the resting period, both PBS-injected groups had an energy intake (50.75 kJ/day and 60.13 kJ/day for resting and activity periods, respectively) that is similar to that required by *C. perspicillata* in the wild to meet daily energy expenditure (50 kJ/day; Speakman and Król 2010). The effect of LPS on energy balance was considerable: mean food intake when bats were injected during the resting period provided 24.56, 28.25 and 14.18 kJ (from the lowest to the highest dose), whereas, when injected during the activity period, mean food intake provided 23.47, 3.80 and 3.05 kJ (from the lowest to the highest dose). Even though bats were caged during the experiment and their energy needs might be lower than free-ranging individuals, it is clear that LPS exposure had dramatic effects on their energy balance in both periods of injections. However, injections of 10 and 15 mg/kg LPS resulted in a more robust decrease in energy intake during the activity period (92.38 – 93.88%) than during the resting period (43.48–71.62%). Rather than considering it as a passive consequence of sickness, decreased food intake has been suggested to be part of an adaptive response to confront disease because it might reduce foraging-related energy expenses and predation risk, as well as lower the availability of micronutrients necessary for pathogens by limiting energy intake, and hence limit the infection (Johnson 2002). In this sense, our results indicate that expression of the anorexic disease behavior is regulated by the strength of the infection during the activity period, with a more robust expression at higher doses, indicating a synergic effect between LPS dose and period of injection.

Body mass loss over a 24-hour period might reflect the mobilization of nutrient stores to cover the energetic cost associated with mounting an immune response and the decrease in food intake by anorexia (Owen-Ashley and Wingfield 2007). Interestingly, dose-dependence in decreased food intake or even a synergic effect of LPS dose and time of injections was not reflected in dose-dependence or synergic effect on mean body mass loss. Mean body mass loss of *C. perspicillata* injected with LPS during the activity and resting periods up to three times (6.4%) than for bats injected with PBS (2%). This change on mean body mass was within the range of changes reported for bats (6-11% decrease after diurnal or nocturnal injection of 1-5 mg/kg LPS; Schneeberger et al. 2013; Stockmaier et al. 2015, 2018; Otálora-Ardila et al. 2016; Cabrera-Martínez et al. 2018; Guerrero-Chacón et al. 2018; Cabrera-Martinez et al. 2019), birds (1-8.4% decrease after diurnal injection of 1 mg/kg LPS; Burness et al., 2010, Owen-Ashley et al. 2008) and small rodents (4.8-9.9% decrease after diurnal injection of 0.05-3.5 mg/kg LPS; Kozak et al. 1994, MacDonald et al. 2011, 2012). In this way, our results indicate that the magnitude of this response may not depend only of dose or period of injection but it may be contingent upon the specific context of the study (e.g., experimental design) and/or a species-specific response. Additionally, despite we did not measure the activity pattern during this study, it is proposed that reduced locomotion due to sickness behavior should ameliorate some costs associated with mounting the immune response (Hart 1988), but there is still an energy deficit (Ashley and Wingfield 2012) which might explain the lack of dose dependence or synergic effect of dose and period of injection on body mass change.

Regardless of the injection time or LPS dose, there was no significative increase in the total leukocyte counts 24 hours after LPS injections. *C. perspicillata* has been previously injected with LPS during the resting period with evidence of leukocytosis (2 mg/kg LPS; Schneeberger et al. 2013), and without evidence of leukocytosis when treated during the activity and resting periods (3 mg/kg LPS; Cabrera-Martinez et al. 2019; Melhado et al. 2020). All previous studies with *C. perspicillata* injected subcutaneously a similar lower LPS dose compared to those used in our study, but Schneeberger et al. (2013) and Melhado et al. (2020) counted WBCs 24 hours after injection whereas Cabrera-Martinez et al. (2019) did it after a shorter period (9 hours). A common feature of the four studies, including the present study, is the existence of individual variation in bat response with some individuals increasing, decreasing or maintaining the number of WBCs after LPS injection. The response of WBCs to LPS has been measured only in three bat families (Molossidae, Phyllostomidae and Pteropodidae) and varying patterns in cellular response to LPS have also been found among species (Stockmaier et al. 2015, 2018; Weise et al. 2017; Guerrero-Chacón et al. 2018). WBC counts increase 8 hours after intraperitoneal injection in *C. plicatus* injected during the resting period (Molossidae; Weise et al. 2017), but no change was found in *M. molossus* 24 hours after subcutaneous injection in the active period (Stockmaier et al. 2015). WBC counts in *D. rotundus* increase 24 hours after subcutaneous injection during the activity period (Phyllostomidae; Stockmaier et al. 2018) but no change was found in *A. lituratus* (Phyllostomidae; Guerrero-Chacón et al. 2018) 24 hours after subcutaneous injection in the resting period. Finally, no change was found in *R. aegyptiacus* 24 hours after subcutaneous injection (Pteropodidae; Moreno et al. 2021). Some of these differences must reflect species-specific differences and/or are contingent on the specific context of the study (e.g., experimental design and response time interval). Further studies with members of other families will allow testing phylogenetic hypotheses.

Similarly to previous studies with bats, LPS doses triggered an increase in N/L when injected during the resting and activity periods (Stockmaier et al. 2018; Cabrera-Martinez et al. 2019; Melhado et al. 2020; Moreno et al. 2021), but this increase was dose-dependent. Dose-dependent increase in N/L was more evident in bats injected with LPS doses during the activity period. Neutrophils constitute the first line of defense against bacterial infections and are the primary phagocytic cells that rapidly proliferate in response to infections and inflammation (Nathan 2006; Davis et al. 2008; Akira et al. 2016). In early stages of infection, neutrophils migrate to affected regions where they are essential in the fight against infections (Shiraishi et al. 1996; Turmelle et al. 2010). LPS activates the HPA axis, culminating in glucocorticoids (GCs) release, promoting the immune cell redistribution (Bornstein et al., 2006; Cain and Cidlowski, 2017). The GCs increase migration of bone marrow-derived neutrophils to the blood stream, while redirect traffic of circulating lymphocytes from the blood stream to lymphoid tissues and bone marrow (Haynes and Fauci 1978; Baschant and Tuckermann 2010; Taves and Ashwell 2021). In this sense, the dose-dependent increase in the N/L ratio may be a result of dose-dependence increase in glucocorticoids in blood stream which consequently redirect neutrophils and lymphocytes traffic (Dhabhar 2002). Additionally, increase in N/L ratio were more robust in bats injected with LPS during the activity period regardless of dose. This robust increase in N/L ratio may be a result of a higher prevalence of glucocorticoids in blood stream during the activity period (Scheiermann et al. 2013; Gong et al. 2015; Markowska et al. 2017) improving immunological function against encounter with bacteria, which are likely to be maximal during the active phase (Scheiermann et al. 2013).

Our results indicate a dose-dependent effect on the duration, but no in the magnitude, of the increase in skin temperature. Body temperature increase peaked between 2 and 4 hours after injections, similarly to previous studies with bats (Otálora-Ardila et al. 2017; Cabrera-Martinez et al. 2019), small rodents (Romanovsky et al. 1996a; Morrow and Opp 2005; Rudaya et al. 2005; MacDonald et al. 2012) and birds (Nomoto 1996; Maloney and Gray 1998; Skold-Chiriac et al. 2015). Additionally, bats injected with the highest dose of LPS maintained a significant increase in skin temperature for a longer time (3 to 5 hours), with a second period of increase in skin temperature at 8 hours (biphasic fever), indicating that this dose activates fever bias longer than lower doses as observed in previous studies with small rodents (Romanovsky et al. 1996a; Rudaya et al. 2005). However, it is important to note that peak of temperature increase after LPS injections (0.5 to 0.9°C) as well as the duration of the febrile response after the highest LPS dose was not higher or lasted longer than observed in previous studies with bats, small rodents and birds that administrated LPS doses at least three times lower (bats: up to 3°C; 7 hours after 1.75-2.80 mg/kg LPS; Otálora-Ardila et al. 2017, Guerrero-Chacón et al. 2018; Cabrera-Martinez et al. 2019; small rodents: up to 3°C 2-11 h after 0.001-1mg/kg LPS; Romanovsky et al. 1996a; Morrow and Oppa, 2005; Rudaya et al. 2005; MacDonald et al. 2012; birds: up to 1.2°C 5-20 hours after 0.001-1mg/kg LPS; Nomoto, 1996; Maloney and Gray 1998; Adelman et al. 2010; Coon et al. 2011; Skold-Chiriac et al. 2015). Based on the limited work available on bats treated with LPS, we speculate that fever is a predominant response, with higher LPS doses being able to trigger long-lasting febrile response, but the magnitude and duration of this response must vary depending on the specie independently of LPS dosage. The functional basis to explain this pattern is not known, but a recent study demonstrates that big brown bats (*Eptesicus fuscus*) have possibly evolved a mechanism to control the over expression of inflammatory genes (TNFα) in response to activation of their innate immune system by viral nucleic acid PAMPS (Banerjee et al. 2017). On other hand, it is possible that, similarly to small rodents and birds, the LPS receptor proteins on the surface of cells (TLR4; Toll-like receptors) is under varying selection, modifying their ability to recognize and respond to LPS (Fornůsková et al. 2013; Vinkler et al. 2014). Subsequent haplotypes of TLRs and their specific ligands can result in altered thermoregulatory responses and contribute to the adaptation of pathogen-host interaction (Fornůsková et al. 2013; Jiang et al. 2017). In fact, recent studies with bats have shown that TLR3, 7, 8 and 9 (viral receptor proteins) sequences in eight bats from three different families (Pteropodidae, Vespertilionidae, and Phyllostomidae) are evolving under purifying selection and multiple mutations in the ligand-binding domain of this receptors have been reported (Romanovsky et al. 1996b; Escalera-Zamudio et al. 2015; Banerjee et al. 2017; Jiang et al. 2017).

## Conclusions

Our study offers important insights into the dependence of time as well as the LPS dosage effect in the APR of bats, and how they deal with the magnitude of infections at different time of day. In general, the results showed that LPS-evoked changes in skin temperature, food intake, and N/L depend on the LPS doses and/or time when LPS is administered (resting or activity period), whereas changes in body mass and WBC must be contingent upon other factors than dose and time of injection. Allied to the fact that the direction of food intake, N/L ratio changes is not determined exactly by the same factors or interaction between factors, and that, apparently thermoregulatory response bats appear to be less sensitive to immune challenge with higher LPS doses, our results suggest that studies that explore the components of APR should consider the LPS doses and time of day in which the response is induced.

## Acknowledgments

We thank Augusto G. Paulino, Gabriel Melhado, Lucia V. Cabrera-Martínez, Pedro Henrique Miguel, Poliana Arantes, Victor H. Bruno, Murilo Negreiros and Ayrton Nascimento, for helping us in the field and/or in the performance of experimental tests.

## Declarations

### Author contributions

A. P. C. N. and L. G. H. M. conceived the ideas and designed methodology. M. F. V. carried out the experiments, collected the data and led the writing of the manuscript. M. F. V. and A. P. C. N. analyzed the data. All authors contributed critically to the drafts and gave final approval for publication.

### Ethics approval

Permits to capture and housing bats were issued by the Instituto Chico Mendes de Conservação da Biodiversidade (ICMBio, process number 66452-1). Ethical permits for this study were issued by the Animal ethics committee of the Universidade Estadual Paulista at Rio Claro (Authorization: n° 3381).

### Funding and Competing interests

This study was supported by a grant to A. P. C. N and L. G. H. M. from Fundação de Amparo à Pesquisa do Estado de São Paulo (FAPESP Visiting Research Program #2017-17607- 6). A. P. C. N. was also was supported by a grant from Fundação de Amparo à Pesquisa do Estado de São Paulo (FAPESP, Grant #2014/16320-7) and L. G. H. M. by a grant from the PASPA-DGAPA program of the Universidad Nacional Autónoma de México (#814-2018). M. F.V. was supported by a grant from the Coordenação de Aperfeiçoamento de Pessoal de Nível Superior Tecnológico (CAPES, Grant #88882.434214/2019-01). The authors declare no conflict of interest.

## Supplementary Material 1 for

**Fig. S1.**
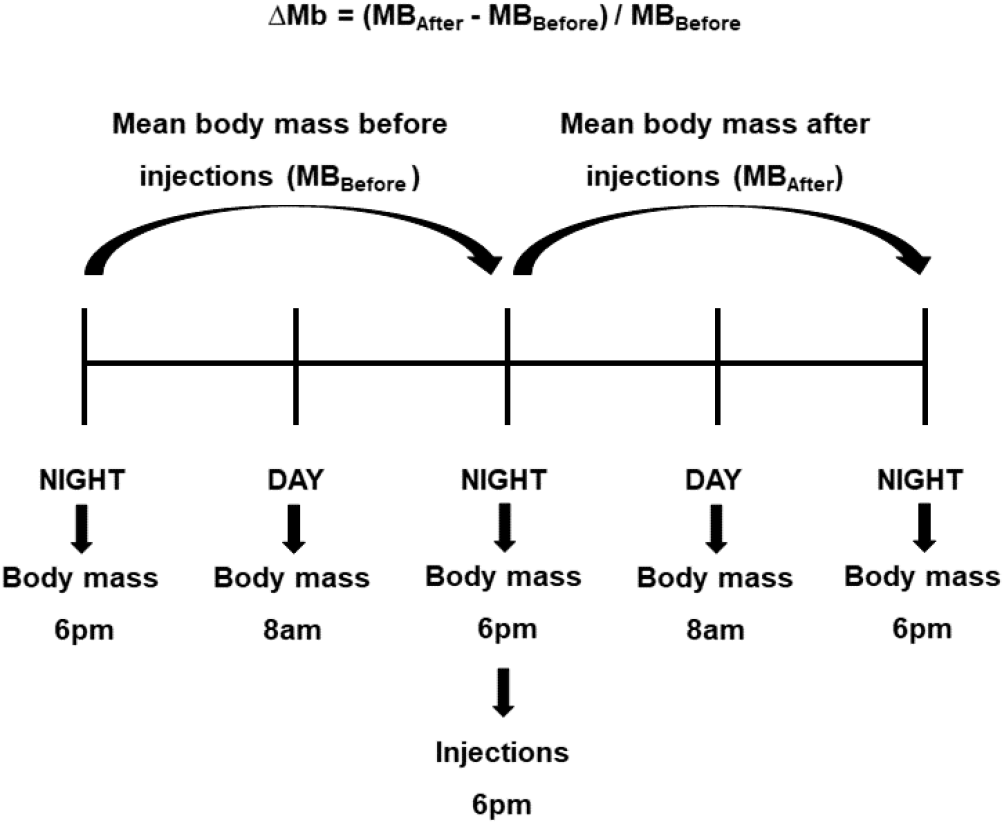
Body mass change (ΔMb) of bats injected during activity period. Body mass changes was assessed in relative terms as: body mass change (ΔMb) = (mean body mass after injections – mean body mass before injections) / (mean body mass before injections)

**Fig. S2.**
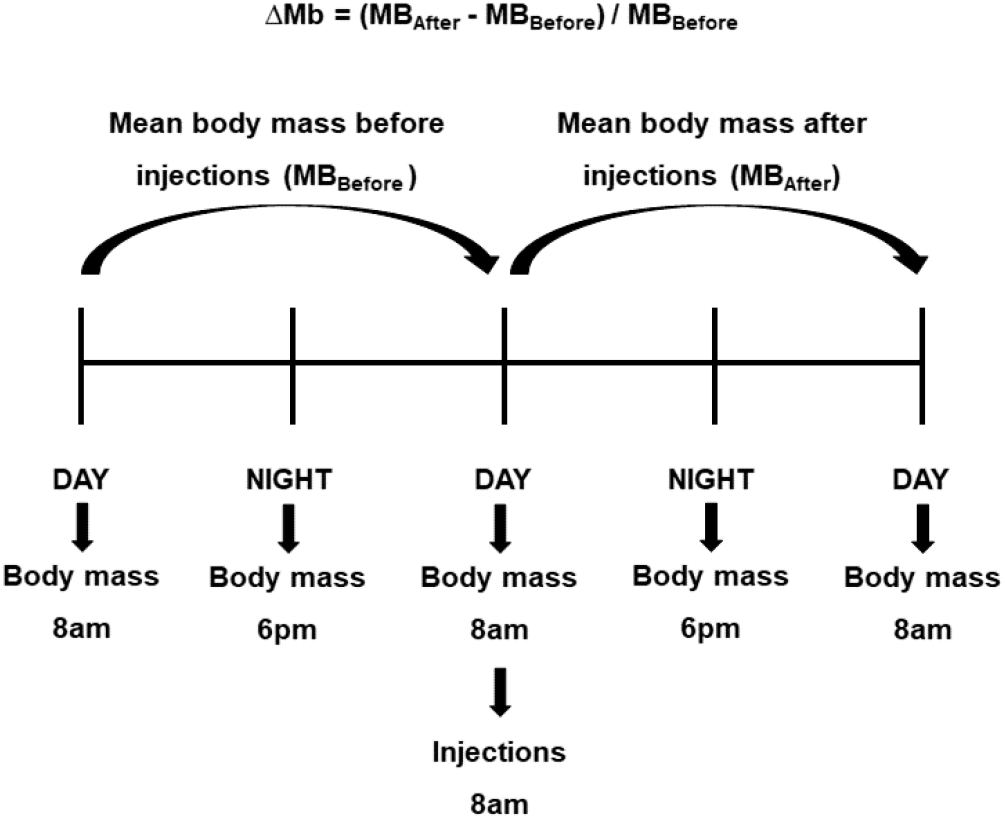
Body mass change in relative terms (ΔMb) of bats injected during resting period. Body mass changes was assessed in relative terms as: body mass change (ΔMb) = (mean body mass after injections – ean body mass before injections) / (mean body mass before injections)

**Fig. S3.**
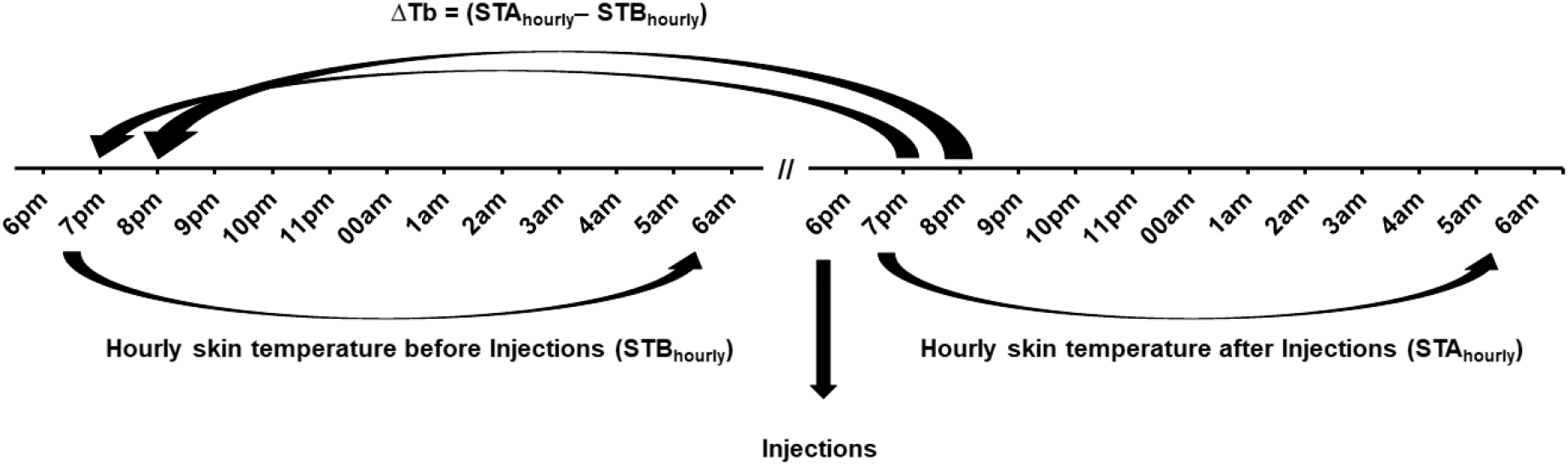
Hourly skin temperature changes in absolute terms (ΔTb) of bats injected during activity period. Hourly skin temperature change (ΔTb) was assessed 12 h after injections in absolute terms by subtracting hourly skin temperature after injections (STA_hourly_) from the respective hourly skin temperature before injections (STB_hourly_)

**Fig. S4.**
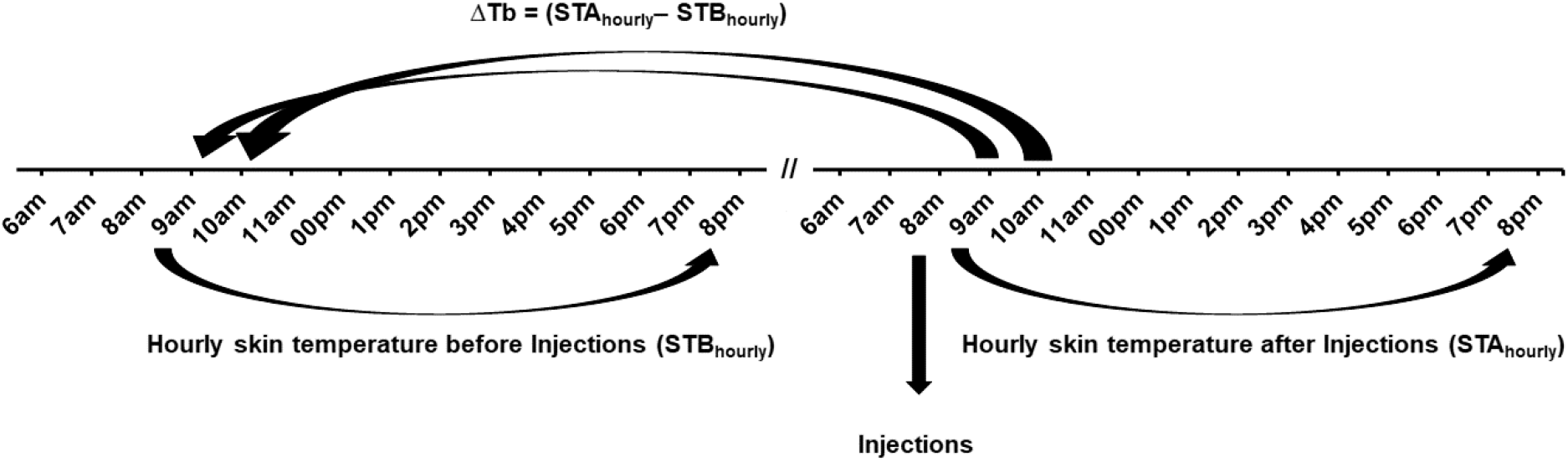
Hourly skin temperature changes in absolute terms (ΔTb) of bats injected during resting period. Hourly skin temperature change (ΔTb) was assessed 12 h after injections in absolute terms by subtracting hourly skin temperature after injections (STA_hourly_) from the respective hourly skin temperature before injections (STB_hourly_)

**Fig. S5.**
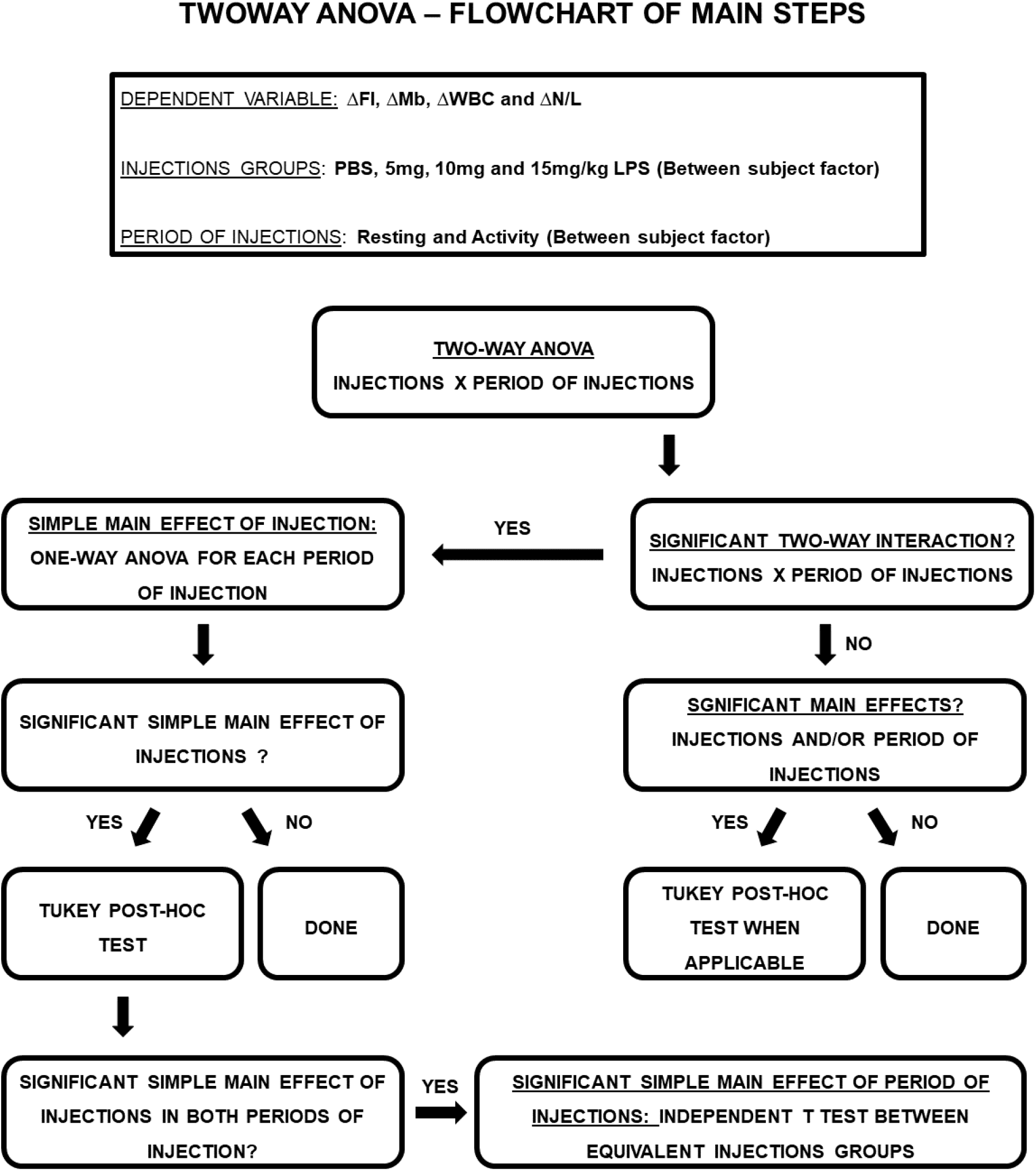
Flowchart of main steps of Two-way ANOVA analysis applied on assessment food intake change (ΔFI), mean body mass change (ΔMb), white blood cell change (ΔWBC) and neutrophil/lymphocyte change (ΔN/L)

**Table S1.**
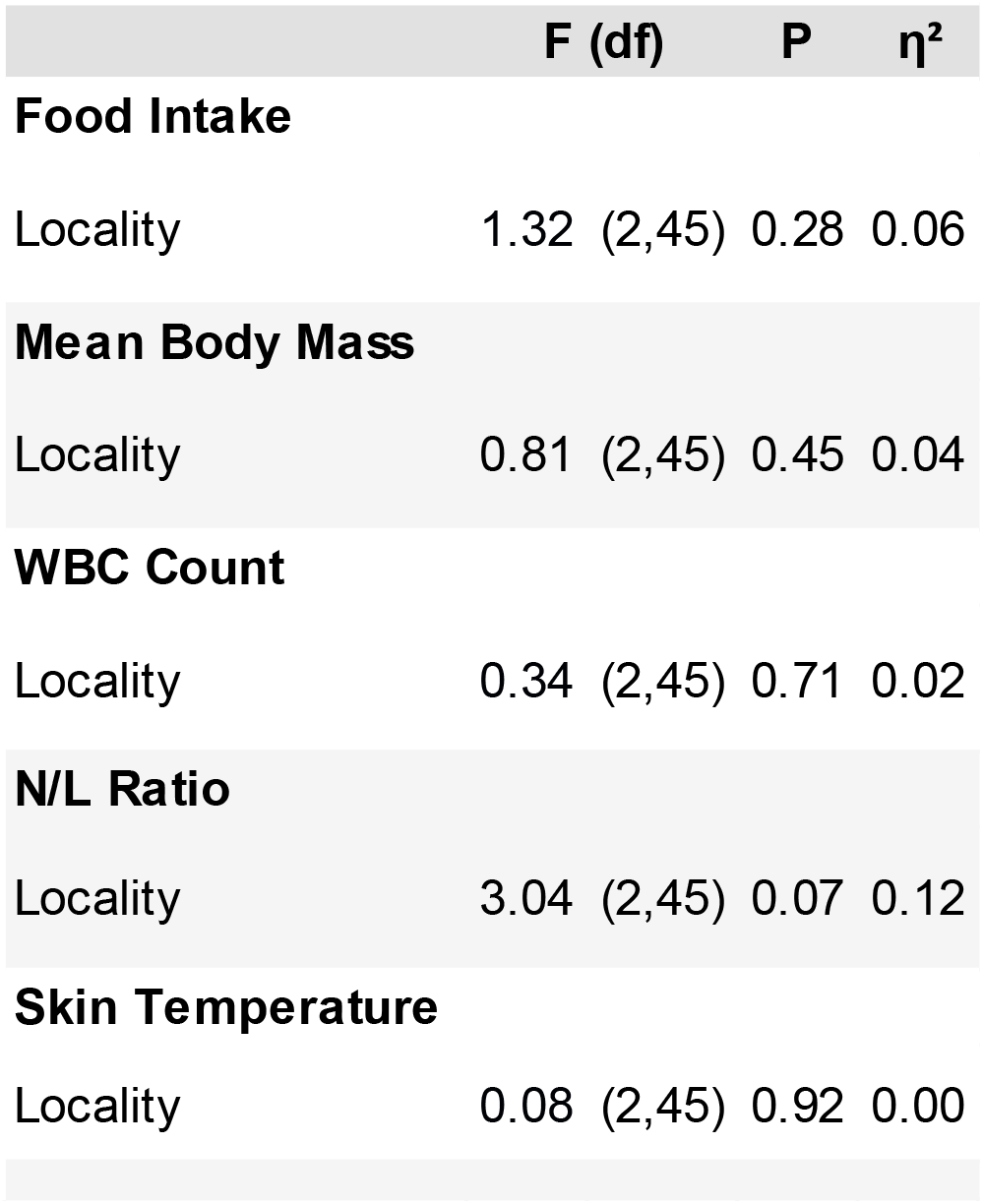
Results of analyses of variance to examine the effect of locality on initial food intake, mean body mass, total white blood cell count (WBC), mean skin temperature and neutrophil/lymphocyte ratio in *Carollia perspicillata*

## Supplementary Material 2 for

**Table S1_1.**
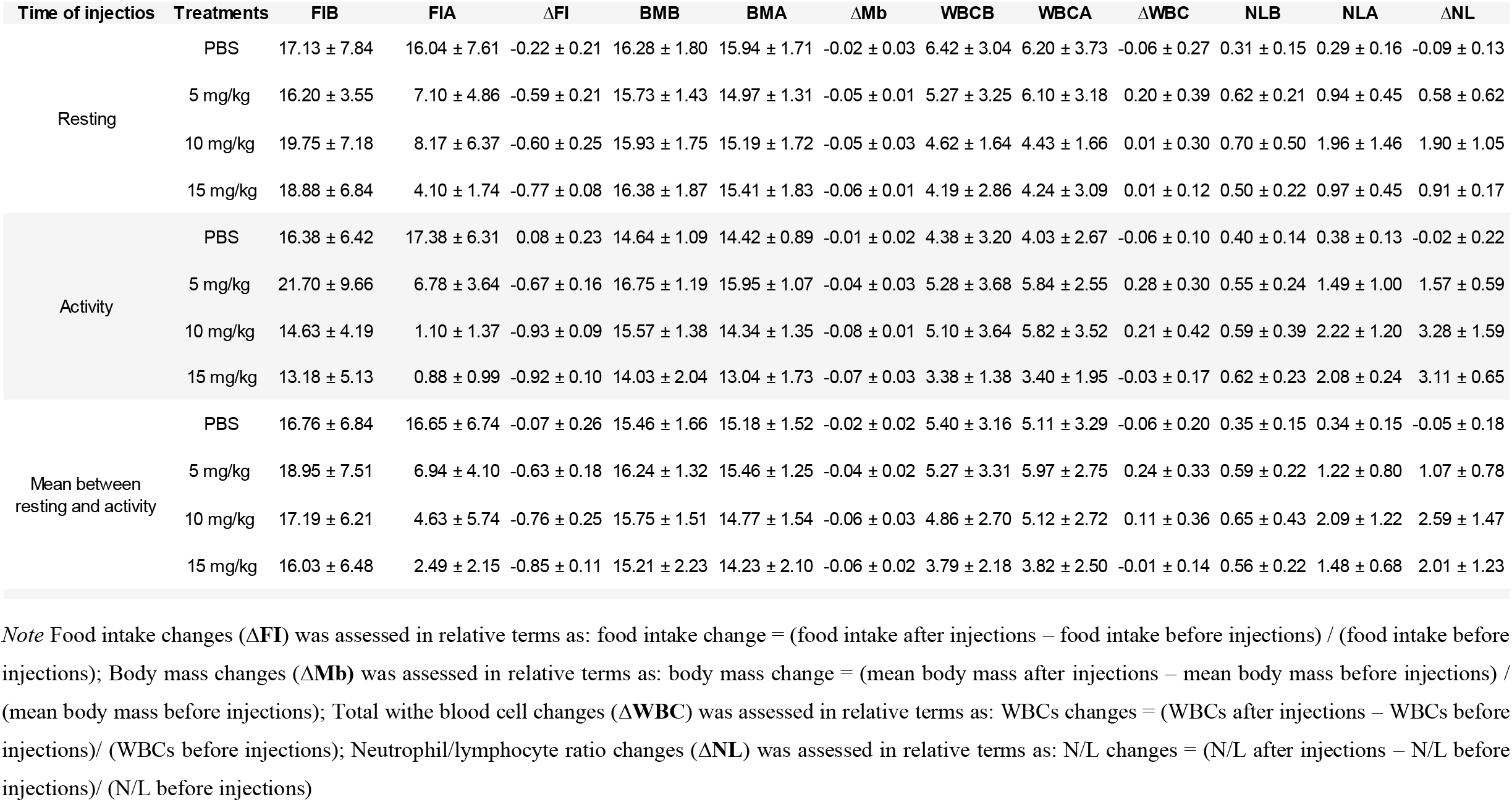
Food intake, body mass, total white blood cell count and neutrophil/lymphocyte ratio before and after PBS and LPS injections (5, 10 and 15 mg/kg LPS) in *Carollia perspicillata*. **FIB**: mean food intake before injections; **FIA**: mean food intake after injections; **ΔFI**: mean food intake changes after injections; **WBCB**: mean total white blood cell count before injections; **WBCA**: mean white blood cell count after injections; **ΔWBC**: mean total white blood cell count change after injections; **NLB**: mean neutrophil/lymphocyte ratio before injections; **NLA**: mean neutrophil/lymphocyte ratio after injections; **ΔNL**: mean neutrophil/lymphocyte ratio change after injections. **BMB**: mean body mass before injections; **BMA**: mean body mass after injections; **ΔMb**: mean body mass change after injections. N = 5 for ΔFI in PBS injected groups. N = 6 for all other variables in PBS and LPS injected groups. ± Standard deviation

**Table S2.**
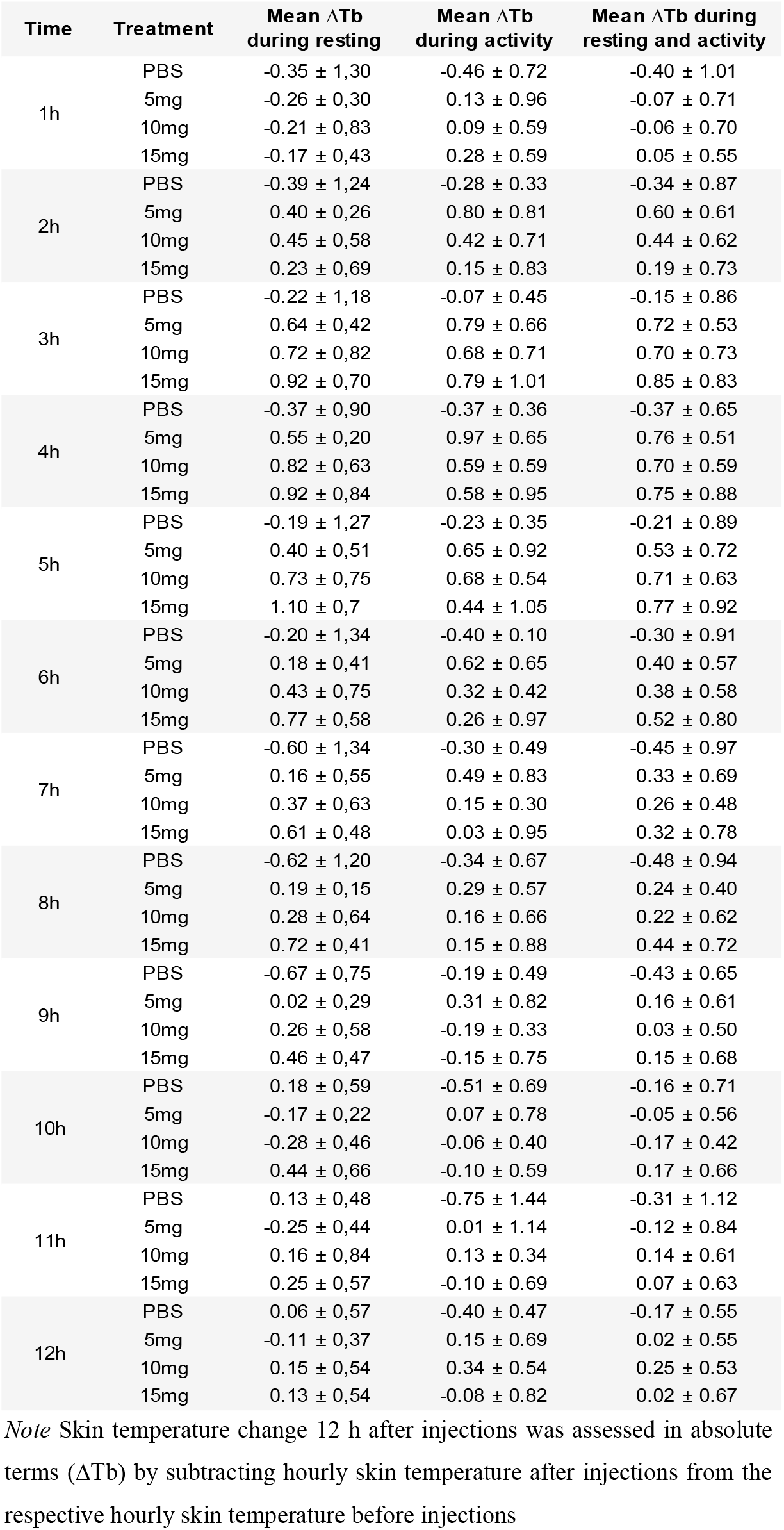
Hourly skin temperature changes (ΔTb) after PBS and LPS injections (5, 10 and 15 mg/kg LPS) during resting and activity periods in *Carollia perspicillata*. N = 6 for all groups. ± Standard deviation

**Table S3.**
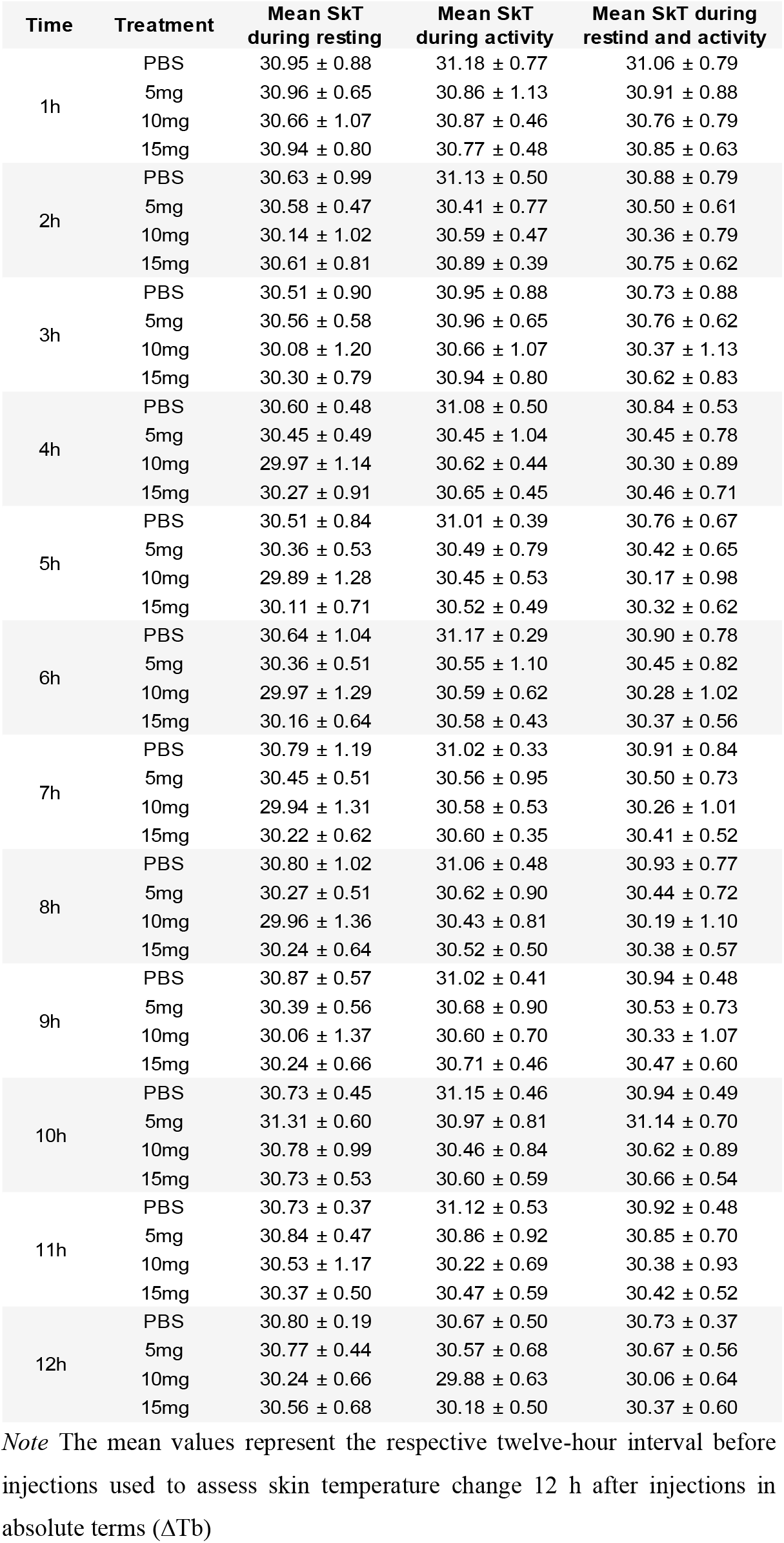
Hourly skin temperature (SkT) before PBS and LPS injections (5, 10 and 15 mg/kg LPS) during resting and activity periods in *Carollia perspicillata*. N = 6 for all groups. ± Standard deviation

**Table S4.**
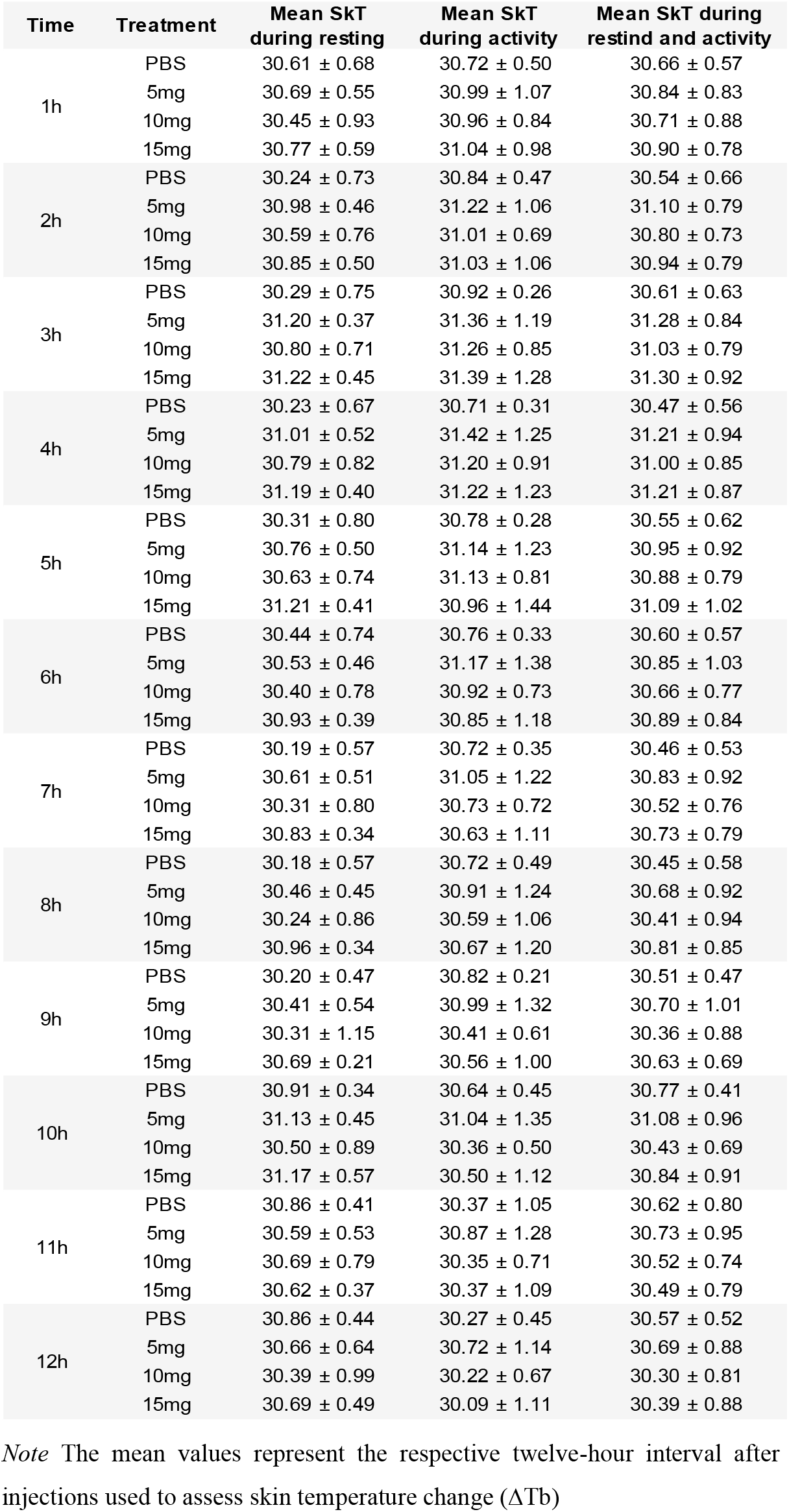
Hourly skin temperature (SkT) after PBS and LPS injections (5, 10 and 15 mg/kg LPS) during resting and activity periods in *Carollia perspicillata*. N = 6 for all groups. ± Standard deviation

## Notes

### Competing Interest Statement

The authors have declared no competing interest.

## References

1. Acevedo-Whitehouse K, Duffus ALJ (2009) Effects of environmental change on wildlife health. Philos Trans R Soc B Biol Sci 364:3429–3438. https://doi.org/10.1098/rstb.2009.0128

2. Adelman JS, Córdoba-Córdoba S, Spoelstra K, et al (2010) Radiotelemetry reveals variation in fever and sickness behaviours with latitude in a free-living passerine. Funct Ecol 24:813–823. https://doi.org/10.1111/j.1365-2435.2010.01702.x

3. Akira S, Lanier L, Nuñes G, et al (2016) Innate Immunity: The First Lines of Defense. In: Murphy K, Weaver C (eds) Janeway’s immunobiology. Garland science, pp 77–138

4. Amaral-Silva L do, Gargaglioni LH, Steiner AA, et al (2021) Regulated hypothermia in response to endotoxin in birds. J Physiol 599:2969–2986. https://doi.org/10.1113/JP281385

5. Amaral-Silva L, Tazawa H, Bícego KC, Burggren WW (2020) Metabolic and Hematological Responses to Endotoxin-Induced Inflammation in Chicks Experiencing Embryonic 2,3,7,8-Tetrachlorodibenzodioxin Exposure. Environ Toxicol Chem 39:2208–2220. https://doi.org/10.1002/etc.4832

6. Ashley NT, Wingfield J (2012) Sickness Behavior in Vertebrates: Allostasis, Life-History Modulation, and Hormonal Regulation. In: Demas GE, Nelson RJ (eds) Ecoimmunology. Oxford: Oxford University Press., pp 45–91

7. Banerjee A, Rapin N, Bollinger T, Misra V (2017) Lack of inflammatory gene expression in bats: A unique role for a transcription repressor. Sci Rep 7:1–15. https://doi.org/10.1038/s41598-017-01513-w

8. Baschant U, Tuckermann J (2010) The role of the glucocorticoid receptor in inflammation and immunity. J Steroid Biochem Mol Biol 120:69–75. https://doi.org/10.1016/j.jsbmb.2010.03.058

9. Basso JC, Shang A, Elman M, et al (2015) Acute Exercise Improves Prefrontal Cortex but not Hippocampal Function in Healthy Adults. J Int Neuropsychol Soc 21:791–801. https://doi.org/10.1017/S135561771500106X

10. Bicego KC, Steiner AA, Antunes-Rodrigues J, Branco LGS (2002) Indomethacin impairs LPS-induced behavioral fever in toads. J Appl Physiol 93:512–516. https://doi.org/10.1152/japplphysiol.00121.2002

11. Burness G, Armstrong C, Fee T, Tilman-Schindel E (2010) Is there an energetic-based trade-off between thermoregulation and the acute phase response in zebra finches? J Exp Biol 213:1386–1394. https://doi.org/10.1242/jeb.027011

12. Cabrera-Martínez L V., Herrera M. LG, Cruz-Neto AP (2018) The energetic cost of mounting an immune response for Pallas’s long-tongued bat (Glossophaga soricina). PeerJ 6: e4627. https://doi.org/10.7717/peerj.4627

13. Cabrera-Martinez L V., Herrera M. LG, Cruz-Neto AP (2019) Food restriction, but not seasonality, modulates the acute phase response of a Neotropical bat. Comp Biochem Physiol Part A Mol Integr Physiol 229:93–100. https://doi.org/10.1016/j.cbpa.2018.12.003

14. Christopoulos P, Kirchner M, Bozorgmehr F, et al (2019) Identification of a highly lethal V3 + TP53 + subset in ALK + lung adenocarcinoma. Int J Cancer 144:190–199. https://doi.org/10.1002/ijc.31893

15. Cohen A, Zemel C, Colodner R, et al (2020) Interactive role of endocrine stress systems and reproductive hormones in the effects of stress on declarative memory. Psychoneuroendocrinology 120:104807. https://doi.org/10.1016/j.psyneuen.2020.104807

16. Coon CAC, Warne RW, Martin LB (2011) Acute-phase responses vary with pathogen identity in house sparrows (Passer domesticus). Am J Physiol Integr Comp Physiol 300:R1418–R1425. https://doi.org/10.1152/ajpregu.00187.2010

17. Copeland S, Warren HS, Lowry SF, et al (2005) Acute Inflammatory Response to Endotoxin in Mice and Humans. Clin Diagnostic Lab Immunol 12:60–67. https://doi.org/10.1128/CDLI.12.1.60-67.2005

18. Cray C, Zaias J, Altman NH (2009) Acute phase response in animals: a review. Comp Med 59:517–26

19. Cruz-Neto, A.P. and Jones K. (2006) Exploring the evolution of the basal metabolic rate in bats. In: Zubaid A, McCracken GFM, Kunz TH (eds) Functional and Evolutionary Ecology of Bats. Oxford University Press, New York, pp 56–89

20. Csabafi K, Bagosi Z, Dobó É, et al (2018) Kisspeptin modulates pain sensitivity of CFLP mice. Peptides 105:21–27. https://doi.org/10.1016/j.peptides.2018.04.018

21. Dantonio V, BatalhÃ O ME, Fernandes MHMR, et al (2016) Nitric oxide and fever: Immune-to-brain signaling vs. thermogenesis in chicks. Am J Physiol - Regul Integr Comp Physiol 310: R896–R905. https://doi.org/10.1152/ajpregu.00453.2015

22. Davis AK, Maney DL, Maerz JC (2008) The use of leukocyte profiles to measure stress in vertebrates: a review for ecologists. Funct Ecol 22:760–772. https://doi.org/10.1111/j.1365-2435.2008.01467.x

23. Demas G.E., Nelson RJ (2012) Introduction to ecoimmunology. In: Demas GE, Nelson RJ (eds) Ecoimmunology. Oxford: Oxford University Press., pp 3–7

24. Demas GE, Zysling DA, Beechler BR, et al (2011) Beyond phytohaemagglutinin: assessing vertebrate immune function across ecological contexts. J Anim Ecol 80:710–730. https://doi.org/10.1111/j.1365-2656.2011.01813.x

25. Dhabhar FS (2002) Stress-induced augmentation of immune function—The role of stress hormones, leukocyte trafficking, and cytokines. Brain Behav Immun 16:785–798. https://doi.org/10.1016/S0889-1591(02)00036-3

26. Do Amaral JPS, Marvin GA, Hutchison VH (2002) The influence of bacterial lipopolysaccharide on the thermoregulation of the box turtle Terrapene carolina. Physiol Biochem Zool 75:273–282. https://doi.org/10.1086/341816

27. Escalera-Zamudio M, Zepeda-Mendoza ML, Loza-Rubio E, et al (2015) The evolution of bat nucleic acid-sensing Toll-like receptors. Mol Ecol 24:5899–5909. https://doi.org/10.1111/mec.13431

28. Fornůsková A, Vinkler M, Pagès M, et al (2013) Contrasted evolutionary histories of two Toll-like receptors (Tlr4 and Tlr7) in wild rodents (MURINAE). BMC Evol Biol 13. https://doi.org/10.1186/1471-2148-13-194

29. Gong S, Miao Y-L, Jiao G-Z, et al (2015) Dynamics and Correlation of Serum Cortisol and Corticosterone under Different Physiological or Stressful Conditions in Mice. PLoS One 10: e0117503. https://doi.org/10.1371/journal.pone.0117503

30. Guan L, Yu WS, Shrestha S, et al (2020) TTC9A deficiency induces estradiol-mediated changes in hippocampus and amygdala neuroplasticity-related gene expressions in female mice. Brain Res Bull 157:162–168. https://doi.org/10.1016/j.brainresbull.2020.02.004

31. Guerrero-Chacón AL, Rivera-Ruíz D, Rojas-Díaz V, et al (2018) Metabolic cost of acute phase response in the frugivorous bat, Artibeus lituratus. Mammal Res 63:397–404. https://doi.org/10.1007/s13364-018-0375-z

32. Haba R, Shintani N, Onaka Y, et al (2012) Lipopolysaccharide affects exploratory behaviors toward novel objects by impairing cognition and/or motivation in mice: Possible role of activation of the central amygdala. Behav Brain Res 228:423–431. https://doi.org/10.1016/j.bbr.2011.12.027

33. Hart BL (1988) Biological Basis of the Behavior of Sick Animals. Vet Clin North Am Small Anim Pract 21:225–237. https://doi.org/10.1016/S0195-5616(91)50028-0

34. Hasselquist D, Nilsson J-Å (2012) Physiological mechanisms mediating costs of immune responses: what can we learn from studies of birds? Anim Behav 83:1303–1312. https://doi.org/10.1016/j.anbehav.2012.03.025

35. Haynes BF, Fauci AS (1978) The differential effect of in vivo hydrocortisone on the kinetics of subpopulations of human peripheral blood thymus-derived lymphocytes. J Clin Invest 61:703–707. https://doi.org/10.1172/JCI108982

36. Jiang H, Li J, Li L, et al (2017) Selective evolution of Toll-like receptors 3, 7, 8, and 9 in bats. Immunogenetics 69:271–285. https://doi.org/10.1007/s00251-016-0966-2

37. Johnson R. (2002) The concept of sickness behavior: a brief chronological account of four key discoveries. Vet Immunol Immunopathol 87:443–450. https://doi.org/10.1016/S0165-2427(02)00069-7

38. Kacprzyk J, Hughes GM, Palsson-McDermott EM, et al (2017) A Potent Anti-Inflammatory Response in Bat Macrophages May Be Linked to Extended Longevity and Viral Tolerance. Acta Chiropterologica 19:219–228. https://doi.org/10.3161/15081109ACC2017.19.2.001

39. Kiernan, D. (2014). Natural Resources Biometrics. Open SUNY Textbooks, Milne Library, State University of New York at Geneseo.

40. Kimura M, Toth LA, Agostini H, et al (1994) Comparison of acute phase responses induced in rabbits by lipopolysaccharide and double-stranded RNA. Am J Physiol Integr Comp Physiol 267: R1596–R1605. https://doi.org/10.1152/ajpregu.1994.267.6.R1596

41. King MO, Swanson DL (2013) Activation of the immune system incurs energetic costs but has no effect on the thermogenic performance of house sparrows during acute cold challenge. J Exp Biol 216:2097–2102. https://doi.org/10.1242/jeb.079574

42. Kozak W, Conn CA, Kluger MJ (1994) Lipopolysaccharide induces fever and depresses locomotor activity in unrestrained mice. Am J Physiol Integr Comp Physiol 266:R125–R135. https://doi.org/10.1152/ajpregu.1994.266.1.R125

43. Krams I, Vrublevska J, Cirule D, et al (2012) Heterophil/lymphocyte ratios predict the magnitude of humoral immune response to a novel antigen in great tits (Parus major). Comp Biochem Physiol Part A Mol Integr Physiol 161:422–428. https://doi.org/10.1016/j.cbpa.2011.12.018

44. Kuzmin I V., Bozick B, Guagliardo SA, et al (2011) Bats, emerging infectious diseases, and the rabies paradigm revisited. Emerg Health Threats J 4:7159. https://doi.org/10.3402/ehtj.v4i0.7159

45. Lee KA (2006) Linking immune defenses and life history at the levels of the individual and the species. Integr Comp Biol 46:1000–1015. https://doi.org/10.1093/icb/icl049

46. Lind CM, Agugliaro J, Farrell TM (2020) The metabolic response to an immune challenge in a viviparous snake, Sistrurus miliarius. J Exp Biol 223:. https://doi.org/10.1242/jeb.225185

47. Liu E, Lewis K, Al-Saffar H, et al (2012) Naturally occurring hypothermia is more advantageous than fever in severe forms of lipopolysaccharide- and Escherichia coli-induced systemic inflammation. Am J Physiol Integr Comp Physiol 302: R1372–R1383. https://doi.org/10.1152/ajpregu.00023.2012

48. Llewellyn D, Brown GP, Thompson MB, Shine R (2011) Behavioral responses to immune-system activation in an anuran (the cane toad, Bufo marinus): Field and laboratory studies. Physiol Biochem Zool 84:77–86. https://doi.org/10.1086/657609

49. Lochmiller RL, Deerenberg C (2000) Trade-offs in evolutionary immunology: just what is the cost of immunity? Oikos 88:87–98. https://doi.org/10.1034/j.1600-0706.2000.880110.x

50. Love AC, Crooks N, Ford AT (2020) The effects of wastewater effluent on multiple behaviours in the amphipod, Gammarus pulex. Environ Pollut 267:115386. https://doi.org/10.1016/j.envpol.2020.115386

51. MacDonald L, Begg D, Weisinger RS, Kent S (2012) Calorie restricted rats do not increase metabolic rate post-LPS, but do seek out warmer ambient temperatures to behaviourally induce a fever. Physiol Behav 107:762–772. https://doi.org/10.1016/j.physbeh.2012.06.009

52. MacDonald L, Radler M, Paolini AG, Kent S (2011) Calorie restriction attenuates LPS-induced sickness behavior and shifts hypothalamic signaling pathways to an anti-inflammatory bias. Am J Physiol Integr Comp Physiol 301: R172–R184. https://doi.org/10.1152/ajpregu.00057.2011

53. Maloney SK, Gray DA (1998) Characteristics of the febrile response in Pekin ducks. J Comp Physiol - B Biochem Syst Environ Physiol 168:177–182. https://doi.org/10.1007/s003600050134

54. Marais M, Maloney SK, Gray DA (2011) The metabolic cost of fever in Pekin ducks. J Therm Biol 36:116–120. https://doi.org/10.1016/j.jtherbio.2010.12.004

55. Markowska M, Majewski PM, Skwarło-Sońta K (2017) Avian biological clock – Immune system relationship. Dev Comp Immunol 66:130–138. https://doi.org/10.1016/j.dci.2016.05.017

56. Martin LB, Weil ZM, Nelson RJ (2007) Fever and sickness behaviour vary among congeneric rodents. Funct Ecol 22:68–77. https://doi.org/10.1111/j.1365-2435.2007.01347.x

57. Mathias S, Schiffelholz T, Linthorst ACE, et al (2000) Diurnal Variations in Lipopolysaccharide-Induced Sleep, Sickness Behavior and Changes in Corticosterone Levels in the Rat. Neuroendocrinology 71:375–385. https://doi.org/10.1159/000054558

58. Maxwell, S. E., Delaney, H. D. (2004). Designing Experiments and Analyzing Data. A Model Comparison Perspective (2nd ed.). Mahwah, NJ: Lawrence Erlbaum Associates.

59. Melhado G, Herrera M. LG, Cruz-Neto AP (2020) Bats respond to simulated bacterial infection during the active phase by reducing food intake. J Exp Zool Part A Ecol Integr Physiol 333:536–542. https://doi.org/10.1002/jez.2399

60. Merlo JL, Cutrera AP, Zenuto RR (2016) Food Restriction Affects Inflammatory Response and Nutritional State in Tuco-tucos (Ctenomys talarum). J Exp Zool Part A Ecol Genet Physiol 325:675–687. https://doi.org/10.1002/jez.2060

61. Moratelli R, Calisher CH (2015) Bats and zoonotic viruses: can we confidently link bats with emerging deadly viruses? Mem Inst Oswaldo Cruz 110:1–22. https://doi.org/10.1590/0074-02760150048

62. Moreno KR, Weinberg M, Harten L, et al (2021) Sick bats stay home alone: fruit bats practice social distancing when faced with an immunological challenge. Ann N Y Acad Sci. https://doi.org/10.1111/nyas.14600

63. Morrow JD, Opp MR (2005) Diurnal variation of lipopolysaccharide-induced alterations in sleep and body temperature of interleukin-6-deficient mice. Brain Behav Immun 19:40–51. https://doi.org/10.1016/j.bbi.2004.04.001

64. Mühldorfer K (2013) Bats and Bacterial Pathogens: A Review. Zoonoses Public Health 60:93–103. https://doi.org/10.1111/j.1863-2378.2012.01536.x

65. Nathan C (2006) Neutrophils and immunity: Challenges and opportunities. Nat Rev Immunol 6:173–182. https://doi.org/10.1038/nri1785

66. Nomoto S (1996) Diurnal variations in fever induced by intravenous LPS injection in pigeons. Pflugers Arch Eur J Physiol 431:987–989. https://doi.org/10.1007/s004240050096

67. O’Shea TJ, Cryan PM, Cunningham AA, et al (2014) Bat Flight and Zoonotic Viruses. Emerg Infect Dis 20:741–745. https://doi.org/10.3201/eid2005.130539

68. Otálora-Ardila A, Herrera M. LG, Flores-Martínez JJ, Welch KC (2016) Metabolic Cost of the Activation of Immune Response in the Fish-Eating Myotis (Myotis vivesi): The Effects of Inflammation and the Acute Phase Response. PLoS One 11:e0164938. https://doi.org/10.1371/journal.pone.0164938

69. Otálora-Ardila A, Herrera M. LG, Flores-Martínez JJ, Welch KC (2017) The effect of short-term food restriction on the metabolic cost of the acute phase response in the fish-eating Myotis (Myotis vivesi). Mamm Biol 82:41–47. https://doi.org/10.1016/j.mambio.2016.11.002

70. Owen-Ashley NT, Hasselquist D, Råberg L, Wingfield JC (2008) Latitudinal variation of immune defense and sickness behavior in the white-crowned sparrow (Zonotrichia leucophrys). Brain Behav Immun 22:614–625. https://doi.org/10.1016/j.bbi.2007.12.005

71. Owen-Ashley NT, Turner M, Hahn TP, Wingfield JC (2006) Hormonal, behavioral, and thermoregulatory responses to bacterial lipopolysaccharide in captive and free-living white-crowned sparrows (Zonotrichia leucophrys gambelii). Horm Behav 49:15–29. https://doi.org/10.1016/j.yhbeh.2005.04.009

72. Owen-Ashley NT, Wingfield JC (2007) Acute phase responses of passerine birds: characterization and seasonal variation. J Ornithol 148:583–591. https://doi.org/10.1007/s10336-007-0197-2

73. Paksuz S, Paksuz EP, Özkan B (2009) White Blood Cell (Wbc) Count of Different Bat (Chiroptera) Species. Trakya University Journal of Natural Sciences 10:55–59. https://dergipark.org.tr/en/pub/trakyafbd/issue/23000/246000

74. Patz JA, Olson SH, Uejio CK, Gibbs HK (2008) Disease Emergence from Global Climate and Land Use Change. Med Clin North Am 92:1473–1491. https://doi.org/10.1016/j.mcna.2008.07.007

75. Rakus K, Ronsmans M, Vanderplasschen A (2017) Behavioral fever in ectothermic vertebrates. Dev Comp Immunol 66:84–91. https://doi.org/10.1016/j.dci.2016.06.027

76. Ramirez-Otarola N, Sarria M, Rivera DS, et al (2018) Ecoimmunology in degus: interplay among diet, immune response, and oxidative stress. J Comp Physiol B 189:143–152. https://doi.org/10.1007/s00360-018-1195-9

77. Romanovsky AA, Kulchitsky VA, Akulich N V., et al (1996a) First and second phases of biphasic fever: Two sequential stages of the sickness syndrome? Am J Physiol - Regul Integr Comp Physiol 271. https://doi.org/10.1152/ajpregu.1996.271.1.r244

78. Romanovsky AA, Shido O, Sakurada S, et al (1996b) Endotoxin shock: Thermoregulatory mechanisms. Am J Physiol - Regul Integr Comp Physiol 270. https://doi.org/10.1152/ajpregu.1996.270.4.r693

79. Roy S, Kumar V, Kumar V, Behera BK (2016) Acute Phase Proteins and their Potential Role as an Indicator for Fish Health and in Diagnosis of Fish Diseases. Protein Pept Lett 24:78–89. https://doi.org/10.2174/0929866524666161121142221

80. Rudaya AY, Steiner AA, Robbins JR, et al (2005) Thermoregulatory responses to lipopolysaccharide in the mouse: dependence on the dose and ambient temperature. Am J Physiol Integr Comp Physiol 289: R1244–R1252. https://doi.org/10.1152/ajpregu.00370.2005

81. Rygula R, Abumaria N, Domenici E, et al (2006) Effects of fluoxetine on behavioral deficits evoked by chronic social stress in rats. Behav Brain Res 174:188–192. https://doi.org/10.1016/j.bbr.2006.07.017

82. Sampath V (2018) Bacterial endotoxin-lipopolysaccharide; structure, function and its role in immunity in vertebrates and invertebrates. Agric Nat Resour 52:115–120. https://doi.org/10.1016/j.anres.2018.08.002

83. Scheiermann C, Kunisaki Y, Frenette PS (2013) Circadian control of the immune system. Nat Rev Immunol 13:190–198. https://doi.org/10.1038/nri3386

84. Scheiermann C, Kunisaki Y, Lucas D, et al (2012) Adrenergic nerves govern circadian leukocyte recruitment to tissues. Immunity 37:290–301. https://doi.org/10.1016/j.immuni.2012.05.021

85. Schneeberger K, Czirjak GA, Voigt CC (2013) Inflammatory challenge increases measures of oxidative stress in a free-ranging, long-lived mammal. J Exp Biol 216:4514–4519. https://doi.org/10.1242/jeb.090837

86. Schoenle LA, Downs CJ, Martin LB (2018) An Introduction to Ecoimmunology. In: Advances in Comparative Immunology. Springer International Publishing, Cham, pp 901–932

87. Shiraishi M, Suzuki K, Abe T, et al (1996) Diurnal variation in neutrophil function. Environ Health Prev Med 1:65–70. https://doi.org/10.1007/BF02931192

88. Skold-Chiriac S, Nord A, Tobler M, et al (2015) Body temperature changes during simulated bacterial infection in a songbird: fever at night and hypothermia during the day. J Exp Biol 218:2961–2969. https://doi.org/10.1242/jeb.122150

89. Speakman JR, Król E (2010) Maximal heat dissipation capacity and hyperthermia risk: neglected key factors in the ecology of endotherms. J Anim Ecol 79. https://doi.org/10.1111/j.1365-2656.2010.01689.x

90. Steiner AA, Hunter JC, Phipps SM, et al (2009) Cyclooxygenase-1 or -2—which one mediates lipopolysaccharide-induced hypothermia? Am J Physiol Integr Comp Physiol 297: R485–R494. https://doi.org/10.1152/ajpregu.91026.2008

91. Stockmaier S, Bolnick DI, Page RA, Carter GG (2018) An immune challenge reduces social grooming in vampire bats. Anim Behav 140:141–149. https://doi.org/10.1016/j.anbehav.2018.04.021

92. Stockmaier S, Dechmann DKN, Page RA, O’Mara MT (2015) No fever and leucocytosis in response to a lipopolysaccharide challenge in an insectivorous bat. Biol Lett 11:20150576. https://doi.org/10.1098/rsbl.2015.0576

93. Sun X, Zu Y, Li X, et al (2021) Corticosterone-induced hippocampal 5.HT responses were muted in depressive-like state. ACS Chem Neurosci 12:845–856. https://doi.org/10.1021/acschemneuro.0c00334

94. Taves MD, Ashwell JD (2021) Glucocorticoids in T cell development, differentiation and function. Nat Rev Immunol 21:233–243. https://doi.org/10.1038/s41577-020-00464-0

95. Titon SCM, Titon B, Muxel SM, et al (2022) Day vs. night variation in the LPS effects on toad’s immunity and endocrine mediators. Comp Biochem Physiol -Part A Mol Integr Physiol 267:111184. https://doi.org/10.1016/j.cbpa.2022.111184

96. Turmelle AS, Ellison JA, Mendonça MT, McCracken GF (2010) Histological assessment of cellular immune response to the phytohemagglutinin skin test in Brazilian free-tailed bats (Tadarida brasiliensis). J Comp Physiol B 180:1155–1164. https://doi.org/10.1007/s00360-010-0486-6

97. Vinkler M, Bainova H, Bryja J (2014) Protein evolution of Toll-like receptors 4, 5 and 7 within Galloanserae birds. Genet Sel Evol 46:1–12. https://doi.org/10.1186/s12711-014-0072-6

98. Volkoff H, Peter RE (2004) Effects of lipopolysaccharide treatment on feeding of goldfish: Role of appetite-regulating peptides. Brain Res 998:139–147. https://doi.org/10.1016/j.brainres.2003.11.011

99. Weber TE, Small BC, Bosworth BG (2005) Lipopolysaccharide regulates myostatin and MyoD independently of an increase in plasma cortisol in channel catfish (Ictalurus punctatus). Domest Anim Endocrinol 28:64–73. https://doi.org/10.1016/j.domaniend.2004.05.005

100. Wei H, Oei TP, Zhou R (2022) Test anxiety impairs inhibitory control processes in a performance evaluation threat situation: Evidence from ERP. Biol Psychol 168:108241. https://doi.org/10.1016/j.biopsycho.2021.108241

101. Weise P, Czirják GA, Lindecke O, et al (2017) Simulated bacterial infection disrupts the circadian fluctuation of immune cells in wrinkle-lipped bats (Chaerephon plicatus). PeerJ 5: e3570. https://doi.org/10.7717/peerj.3570

102. White RJ, Razgour O (2020) Emerging zoonotic diseases originating in mammals: a systematic review of effects of anthropogenic land-use change. Mamm Rev 50:336–352. https://doi.org/10.1111/mam.12201

103. Williams J, Lambert AM, Long R, Saltonstall K (2019) Does hybrid Phragmites australis differ from native and introduced lineages in reproductive, genetic, and morphological traits? Am J Bot 106:29–41. https://doi.org/10.1002/ajb2.1217

104. Williams JB, Tieleman BI, Shobrak M (2009) Validation of Temperature-Sensitive Radio Transmitters for Measurement of Body Temperature in Small Animals. Ardea 97:120–124. https://doi.org/10.5253/078.097.0115

105. Wong S, Lau S, Woo P, Yuen K-Y (2007) Bats as a continuing source of emerging infections in humans. Rev Med Virol 17:67–91. https://doi.org/10.1002/rmv.520

